# The dynamics of “silent” variation in *Mimulus guttatus*: Codon usage bias and linked selection

**DOI:** 10.64898/2026.05.18.725996

**Authors:** Luis Javier Madrigal-Roca, John K. Kelly

## Abstract

1. Synonymous nucleotide variation, which is remarkably high in *Mimulus guttatus*, can be affected by both codon usage selection (translational efficiency) and linked selection (hitchhiking effects).
2. Codon usage reflects a genome-wide tug-of-war between mutational pressure toward A/T-ending codons and weak selection favoring G/C-ending codons. The outcome is determined largely by gene expression level and localized variation in recombination rate.
3. Using both mechanistic (ROC-SEMPPR) and population genetic models, we find that most genes are weakly selected for codon usage, about 76% yielding scaled selection coefficients (S = 4N_e_s) in the range of 0 to 1. Additionally, 4029 genes, primarily involved in photosynthesis, translation, defense, and phosphate scavenging, experience strong selection (S > 1).
4. Levels of nucleotide variation within genes indicate a strong effect of linked selection. Non-synonymous polymorphism declines in genes with strong purifying selection, and as the rate of (intra-genic) recombination declines. Levels of synonymous polymorphism usually track non-synonymous (owing to background selection), except in genes under the strongest translational selection.
5. Counterintuitively, we find that codon usage selection has a generally positive effect on synonymous nucleotide diversity at 4-fold degenerate positions. Since mutation strongly disfavors the optimal base in *M. guttatus*, codon selection in the range of 0 < S < 2 evens the balance (between selection and mutation) and thus inflates heterozygosity.

## INTRODUCTION

One of the main characteristics of the genetic code is its degeneracy (redundancy) (Crick et al., 1961; Deng et al., 2020; Parvathy et al., 2022). Of the 64 possible codon triplets, three encode stop signals, two specify unique amino acids (tryptophan and the start codon methionine), and the remaining 59 encode 18 amino acids with varying degrees of degeneracy. This degeneracy, where one amino acid is specified by anywhere from two to six distinct codons, has proven a boon for evolutionary biologists. Assuming that synonymous mutations can be treated as selectively neutral, patterns of variation can be used to infer populations sizes, historical migration patterns, demographic events such as population bottlenecks and range expansions, and provide time estimates for the divergence of species. Having a collection of “known neutral sites” also provides a natural internal control to test for the effect of selection on putatively non-neutral sites (Haasl & Payseur, 2016; McDonald & Kreitman, 1991; Nielsen, 2001). Unfortunately, these inferences can be undermined if synonymous mutations are not selectively neutral owing to Codon Usage Selection (CUS; Akashi, 1994; Ikemura, 1981) or if synonymous variation is strongly influenced by selection at linked sites (Begun & Aquadro, 1992; Corbett-Detig et al., 2015).

Codon usage bias (CUB) refers to the empirical pattern where the synonymous codons within an amino acid are used with unequal frequency, biases that vary markedly not only between species but also among genes within the same organism (Galtier et al., 2018; Machado et al., 2020; Parvathy et al., 2022). CUB can be driven by simple mutational biases, but selection is possible when synonymous nucleotide changes alter mRNA structure, translation efficiency, or gene expression levels (Parvathy et al., 2022). CUS can be quite strong as illustrated by examples where synonymous substitutions cause measurable effects such as altered expression of the alcohol dehydrogenase gene in Drosophila (Carlini, 2004). In humans, changes in codon composition in the *KRAS* gene have been associated with modifications in protein expression and tumorigenicity (Lampson et al., 2013). When selection is non-negligible, CUB will reflect the interplay of three evolutionary forces: mutation, natural selection, and genetic drift (Akashi, 1994; Cope & Shah, 2022; Gouy & Gautier, 1982; Hershberg & Petrov, 2008; Z. Zhang et al., 2012).

Here, we apply the ROC-SEMPPR model (Gilchrist et al., 2015) to test for selection on synonymous variation. Incorporating gene expression data, ROC-SEMPPR estimates the ‘Ribosome Overhead Cost’ (ROC) of suboptimal codons. The ribosome can stall on a slowly-decoded codon making it unavailable for translating other mRNAs, incurring a fitness penalty (Landerer et al., 2018). The strength of selection is estimated as *N*_*e*_ *s*, the product of the selection coefficient and the effective population size (Bulmer, 1991; Cope & Shah, 2022; S. Wright, 1931). When selection is strong relative to drift (|*N*_*e*_ *s*| > 1), it will drive the frequency of optimal codons upward, and if strong enough, deplete standing variation at synonymous sites. Conversely, when |*N*_*e*_ *s*| < 1, populations confront a regime (Lynch, 2010; Lynch et al., 2016; McCandlish & Plotkin, 2016) where random genetic drift dominates selection and codon frequencies will reflect the underlying mutational spectrum. McVean & Charlesworth (1999) described an interesting case where diversity can be highest in the transition from weak to strong codon selection (the “diversity hump”). When the mutation rate away from the favored base is much higher than toward the favored base, selection of intermediate strength can inflate synonymous polymorphism relative to the neutral expectation because the two opposing forces are evenly balanced.

We predict that nucleotide diversity at synonymous sites will depend not only on the expression level of the gene, but also the distance of the codon from the gene start site. In many plants, including *M. guttatus*, cross-over events occur most frequently near the 5’ end of genes (around transcription start sites) and decline in frequency toward the middle of genes (Hellsten et al., 2013; Hsu et al., 2022; Okagaki et al., 2018; Wijnker et al., 2013). Codon bias should be stronger near gene starts because selection is less hindered by Hill-Robertson Interference (Hill & Robertson, 1966) among linked codons. Perhaps more importantly, polymorphism at synonymous positions will be less affected by deleterious and adaptive mutations at nearby non-synonymous sites (Castellano et al., 2016; Cutter & Payseur, 2013). Linked selection was first demonstrated by studies showing reduce polymorphism in low-recombination portions of the genome relative to high-recombination portions (Begun and Aquadro 1994), but a similar pattern can emerge for synonymous polymorphism within genes if the gradient in recombination rate is sufficiently strong (Comeron et al., 2008; Hollister et al., 2010). As described above, gene expression level can determine the strength of codon-usage selection, but it can also influence the importance of linked selection. If selection on non-synonymous changes is stronger or more frequent in highly expressed genes, the amount of neutral variation will decline with expression level.

In this paper, we synthesize recently developed genomic and transcriptomic resources from *Mimulus guttatus* and reanalyze the extensive polymorphism data for this species to test population genetic predictions regarding synonymous sites variation. Plant research on CUB has focused on two species from the Rosid clade of dicots (*Arabidopsis thaliana* and *Populus trichocarpa*), the moss *Physcomitrella patens* (Parvathy et al., 2022), and the two primary monocot crop species, maize (Campbell & Gowri, 1990) and rice (Wang & Hickey, 2007). We find that *M. guttatus*, an Asterid, produces qualitatively different patterns of CUB in terms of the optimal base at degenerate positions. We find that the distribution of gene-level selection intensities is strongly right-skewed: CUS affects many genes but is dominant for only a minor fraction of highly expressed genes. Because the efficacy of selection on synonymous sites depends not only on *N*_*e*_ *s* but also on the local recombination environment, optimal codon enrichment peaks near the 5’ end of genes where recombination rates are highest. *M. guttatus* exhibits an intra-genic version of the classic linked selection pattern: Synonymous diversity is highest where recombination is highest. Together, these predictions connect the mechanistic models of codon usage evolution directly to observable patterns in sequence variation and expression data, providing a multi-layered view of silent site evolution.

## MATERIALS AND METHODS

### Data

Because the six-fold degenerate amino acids (Leu, Ser, and Arg) contain both two-fold and four-fold redundant codon sub-families that can experience qualitatively different selective pressures on codon usage (X. Sun et al., 2013). Then, we used the coding sequences from *Mimulus guttatus* var. IM767 v2.1 genome (https://phytozome-next.jgi.doe.gov/info/Mguttatusvar_IM767_v2_1) to count the frequency of codons. We then used the raw counts to calculate a myriad of derived measures that can be used as proxy for understanding the patterns of CUB. We only analyzed genes where the following criteria were met. We verified that each gene began with the canonical AUG start codon, terminated with a recognized stop codon (UGA, UAG, or UAA), and contained no internal premature stop codons.

Lovell et al. (2025) collected gene expression data from two different genotypes of *M. guttatus* (IM62 and IM767). A total of 29 RNA libraries were generated from samples encompassing nine distinct tissue types: vegetative (leaf, root, stem, seedling), reproductive (bud, flower, ovary, pollen), and whole-plant. Leaf tissue was the most deeply sampled, with four independent biological replicates for both IM62 and IM767. Reproductive tissues (bud, flower, ovary, and stem) were sampled in duplicate for IM62 and as singletons for IM767. Pollen and whole seedlings were only collected for IM767. A complete breakdown of biological replicates per tissue and line is provided in Supplementary Table S4.

We remapped the raw RNAseq data to the IM767 genome assembly using the custom pipeline SalmonStreamer (Madrigal-Roca et al., 2025; Patro et al., 2017). SalmonStreamer produces a multisource expression profile with read counts per million (CPM) for each transcript. For each expression source, we calculated the decimal logarithm of the CPM, and then we obtained for each gene the maximum value across all replicates and the mean value. Additionally, we calculated the expression breadth, defined as the number of tissue/line instances where expression expressed in CPM was greater than 1. This convention was maintained in all correlational analysis performed. The expression levels across all samples were used as input for the ROC-SEMPPR model, which combines them to estimate an average protein synthesis rate (φ)—a composite metric that reflects long-term translational demand rather than direct measurements of protein abundance.

The third major set of data for our analyses is DNA sequence polymorphism estimated from whole-genome sequences of 187 highly homozygous inbred lines of *Mimulus guttatus* (syn. *Erythranthe guttata*). These lines originated from a single, predominantly outcrossing natural population at Iron Mountain, Oregon, and were established via single-seed descent for a minimum of eight generations to minimize heterozygosity (Troth et al., 2018). Raw paired-end reads from the 187 highly homozygous inbred lines were mapped to the Mimulus guttatus IM767 v2.1 reference assembly using the BWA-MEM algorithm (Li & Durbin, 2009) with default parameters. The resulting alignments were converted to BAM format, coordinate-sorted, and indexed using SAMtools (Danecek et al., 2021). To mitigate PCR amplification biases, duplicate reads were identified and marked using Picard MarkDuplicates (“Picard Toolkit,” 2019).

Variant calling was performed using the mpileup and call commands within BCFtools (Danecek et al., 2021). Because our downstream population genetic analyses required a complete denominator of callable sites, the model was explicitly parameterized to output both variant and invariant sites across the genome. The raw calls were subsequently subjected to rigorous quality control filtering to ensure high-confidence site sets. We restricted our dataset to biallelic single nucleotide polymorphisms (SNPs) and monomorphic sites. To ensure accurate genotyping within these lines, we required that for any homozygous call, the read count of the major allele be at least five times greater than that of the minor allele. Additionally, sites were required to have a minimum mapping quality (MAPQ) of 30 and a minimum base quality of 20. Finally, sites with more than 20% missing genotypes across all 187 lines were removed.

### CUB metrics

To characterize patterns of codon usage bias (CUB) across the *Mimulus guttatus* reference genome, we calculated several standard metrics for each gene. We quantified the relative preference for individual codons using Relative Synonymous Codon Usage (**RSCU**), which compares a codon’s observed frequency to its expected frequency under uniform synonymous usage (Sharp et al., 1986). To assess the overall magnitude of bias, we calculated the Effective Number of Codons (**ENC**), where lower values approach 20 to indicate extreme bias and values approaching 61 indicate random usage (F. Wright, 1990). We also determined position-specific GC content—specifically at the first (**GC1**), first and second combined (**GC12**), and third positions (**GC3**)—to capture localized base composition variations. Additionally, we measured GC content at synonymous third positions (**GC3s**), which excludes the single-codon amino acids tryptophan and methionine, to better isolate mutational signatures. To account for background nucleotide composition, we computed the Codon Deviation Coefficient, **CDC** (Z. Zhang et al., 2012). CDC measures how far a gene’s actual codon usage departs from what its nucleotide composition alone would predict; higher values indicate stronger bias. Using the measures derived from codon counts, we performed some routinely used exploratory visuals to describe the overall pattern of CUB, including the well-known neutrality, parity, and ENC plots. These metrics are based on the reference genome of *Mimulus guttatus*.

Additionally, we performed G-based tests to formally search for a genome-wide signature of CUB (G_total_), for genes that are deviant from the equal usage expectation (G_gene_), for family of amino acids that biased in terms of codon usage (G_aa_), and finally, for heterogeneity across genes (G_het_). The gene level test is complemented with the intrinsic framework for the CDC calculation (Z. Zhang et al., 2012), that allows the incorporation of compositional bias (purine and GC-content) per gene and test statistically the deviations from expectations using bootstrapping.

Another metric widely used in codon usage bias studies is the Codon Adaptation Index, **CAI** (Sharp & Li, 1987). This metric quantifies the relative adaptiveness of each codon relative to the most frequent codon per amino acid in a reference gene of highly expressed genes, and by doing so, globally assign a value per gene, which is usually interpreted as the effectiveness of selection in codon usage patterns (Sharp & Li, 1987). To define a reference set of genes we identified genes in the *Mimulus guttatus* var. IM767 v2.1 reference genome corresponding cytosolic ribosomes and elongation factors, which are constitutively highly expressed genes. Given that annotation hits are putative we filtered the initial list of candidate genes by preserving highly expressed genes according to our expression data, resulting in a list of 129 highly expressed genes. As a complementary analysis to CAI, we performed an enrichment analysis by comparing the usage frequency of each codon across groups of highly, moderately, and lowly expressed genes. This approach provides an alternative perspective to CAI, which can be influenced by reference bias, as the most frequently used codon in a reference gene for a given amino acid may be strongly affected by mutational forces.

### Exploratory associative analysis

We evaluated the relationship between CDC (a metric that quantifies the deviation of a gene’s codon usage from a null expectation based on its nucleotide composition) and gene expression levels, controlling for gene length, which have been found as a typical confounder (Khandia et al., 2022). Metrics like CDC and ENC are sensitive to sequence length; shorter genes possess fewer codons (smaller sample sizes), leading to stochastic deviations from the expected codon distribution. This random noise creates a statistical artifact where short genes appear to have high bias (high CDC). Given that biological dependencies between expression and evolutionary rates are frequently non-linear (e.g., threshold effects of selection), and that length-bias artifacts follow complex decay distributions, we employed Generalized Additive Models (GAMs). This approach avoids the restrictive assumptions of linearity inherent in standard regression, allowing for a data-driven estimation of the relationship shape. We used the mgcv package in R (Wood, 2004, 2011; Wood et al., 2016). As CDC is bounded between zero and one we utilized a beta error distribution with a logit link function.

To further characterize the relationship between expression and codon bias without the confounding influence of gene length, we computed a length-corrected CDC metric. This was derived by extracting the residuals from a GAM that modeled CDC solely as a function of coding sequence length, *CDCs*(*CDS*_*length*_), where *s* refers to a smooth function (beta error distribution, logit link function). A value of zero indicates a gene has the expected CDC for its length; positive values indicate higher-than-expected bias. Using this corrected metric, we stratified genes into three expression regimes: Low (Bottom 5%), Intermediate (Middle 90%), and High (Top 5%) based on the maximum expression value in log10 scale across tissue sources (Max_Log10_Exp). Differences in length-corrected CDC distributions among these groups were assessed using a Kruskal-Wallis test, followed by a post-hoc Dunn’s test with Benjamini-Hochberg correction for multiple comparisons. Those procedures were performed using the R functions kruskal.test (R Core Team, 2024) and dunn.test::dunn.test (Dinno, 2024), respectively.

### Correspondence analysis

We then performed a correspondence analysis (CA) on the codon count contingency table (codon counts by gene), following (Perriere, 2002). This analysis allowed us to explore the ordination of genes in a multidimensional space defined by codon counts, with a particular focus on whether gene categories based on expression could be differentiated along the main axes. Since CA is agnostic to gene length, we also used RSCU values to project genes into a low-dimensional space defined by principal components. To formally test for differentiation among defined categories in these spaces, we used Multivariate Analysis of Variance (MANOVA) on both CA-derived and PC-derived axes.

### ROC-SEMPPR modeling

Disentangling the effects of mutation, selection, and drift on codon usage presents significant statistical challenges. To simplify modeling the interplay between mutation and selection, researchers often assume a constant mutation pressure across the genome (Cope & Shah, 2022). This assumption underlies mechanistic models such as FMutSel (Yang & Nielsen, 2008)) and ROC-SEMPPR (Gilchrist et al., 2015). Notably, ROC-SEMPPR links codon usage to a measurable physiological hecost: when a ribosome stalls on a slowly-decoded codon, it is unavailable for translating other mRNAs, a penalty termed the ‘Ribosome Overhead Cost’ (ROC). The model treats each codon’s contribution to this cost as the target of selection and enables the simultaneous estimation of gene expression (φ) and selection coefficients (Δη) from sequence data alone or in combination with empirical gene expression measurements (Landerer et al., 2018). Some studies suggest that selection on synonymous sites can be quite strong (Machado et al., 2020) which is illustrated by examples with large phenotypic effects caused by substitution of preferred for unpreferred codons, e.g. altered expression patterns of alcohol dehydrogenase (*Adh*) in Drosophila (Carlini, 2004). In humans, changes in codon composition in the *KRAS* gene have been associated with modifications in protein expression and tumorigenicity (Lampson et al., 2013).

To explicitly model the interplay between mutation and selection shaping CUB in our system, we used the R package AnaCoDa (Landerer et al., 2018). The ROC-SEMPPR model in AnaCoDa applies a mutation–selection framework that incorporates the Ribosome Overhead Cost (ROC) to infer gene- and codon-level parameters from codon counts. This model links codon usage to protein production rates and selection on translational efficiency and accuracy, treating observed codon usage as the result of a balance between mutation bias and selection, with both codon ‘costs’ and gene-specific protein production estimated within a Bayesian hierarchical MCMC framework (Gilchrist et al., 2015). The model assumes that synonymous codons differ in their translational costs (such as elongation time and error rates), and that these costs become more significant as the amount of protein produced by a gene increases. Consequently, selection on codon usage is stronger in highly expressed genes, where inefficient translation incurs greater resource costs. Mutation drives codon usage toward a mutational equilibrium determined by base composition and mutation biases, while selection for translational efficiency and accuracy favors preferred codons. The ROC-SEMPPR model infers both mutational and selective components from genome-wide codon counts (Gilchrist, 2007).

Under this model, at equilibrium, the probability of seeing a synonymous codon *i* from an amino acid group of *n*_*a*_ synonymous codons follows a multinomial logistic distribution:

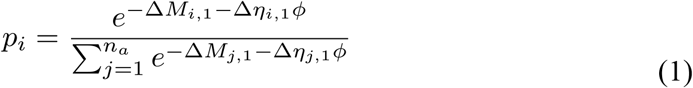

where *p*_*i*_ is the frequency of codon *i* for a given protein synthesis level (*ϕ*), Δ*M*_*i*_,_1_ is the mutational deviation of codon *i* relative to a reference codon in the amino acid family, and *η*_*i*_,_1_is the cost of codon *I* relative to the reference codon (target of selection). From the equation, we can see how a lowly expressed gene is essentially invisible to selection, because no matter the cost, mutation will prevail. So, in essence, we can summarize that

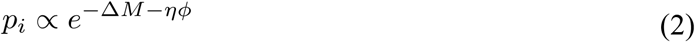

Although the model can simultaneously estimate mutational bias, cost (as the selective target), and protein synthesis parameters, AnaCoDa also allows the incorporation of empirical mutational biases and gene expression data. This feature is particularly useful in complex genomes, where simultaneous estimation of all parameters can lead to identifiability issues.

### Introns and intergenic DNA

We estimated the background mutation parameters (Δ*M*) of ROC-SEMPPR using neutral genomic regions as a proxy. We used intronic sequence trimmed of constrained sites to estimate mutational parameters but also analyzed intergenic flanking regions as a comparison. Sequence Extraction and Filtering Genomic coordinates were obtained from the *Mimulus guttatus* v2.1 annotation (Phytozome). Introns from primary transcripts on main chromosomes were trimmed by 30 bp at both the 5’ and 3’ ends to remove splice site signals. Introns shorter than 86 bp (post-trimming) were discarded. By imposing these criteria, we limit the impact of including potential regulatory elements associated with the spliceosome activity (Machado et al., 2020). For intergenic regions, we extracted non-overlapping sequences 10 kb upstream and downstream of genes, trimming the 1 kb proximal to each gene body to exclude core promoter and terminator elements that experience regulatory constraint. Repetitive elements were excluded by using the hard-masked reference genome (*Mguttatusvar_IM767_887_v2*.*0*.*hardmasked*.*fa*), which was pre-processed to mask transposons and tandem repeats. We calculated nucleotide frequencies (*A*_*freq*_, *C*_*freq*,_ *G*_*freq*,_ *T*_*freq*_) for the extracted sequences (both intronic and intergenic).

Intronic sequences, and not intergenic DNA, was used as a neutral reference to control for local mutation rates and background selection (*N*_*e*_). We chose intronic nucleotide frequencies (T > A > G > C; Table 1) as the mutational baseline because introns are transcribed and reside in open chromatin. Thus, their mutational profile captures any transcription-coupled damage and repair biases that affect coding sequences (Polak & Arndt, 2008) (Niehrs & Luke, 2020; Su & Freudenreich, 2017) (Georgakopoulos-Soares et al., 2020), while being minimally confounded by protein-level selection. We summarized the mutational bias per amino acid by estimating the expected frequency of each codon given stationary nucleotide frequencies, and expressed such mutation bias as log odd ratios of each codon relative to a reference one (last alphabetically to match AnaCoDa convention). For each of the five models, we ran six independent MCMC chains. Convergence was assessed using the Gelman-Rubin statistic and the Geweke diagnostic. Upon confirmation of convergence, we checked whether the *Φ* value was positively correlated with empirical mean expression value (we discarded models where convergence was achieved but correlation was negative). Finally, we extracted posterior estimates for the selection coefficients from the best model. In the ROC-SEMPPR framework, the selection coefficient (*S*) for codon in gene is defined as the product of the intrinsic cost of the codon (Δ *η*_*i*_) and the protein synthesis rate of the gene (Φ_*g*_).

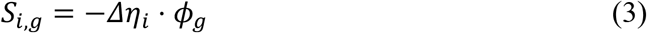

**Table 1:** Observed nucleotide frequencies are reported for Introns and Intergenic regions.

| Region | $A_{freq}$ | $C_{freq}$ | $G_{freq}$ | $T_{freq}$ |
| --- | --- | --- | --- | --- |
| Introns | 0.29 | 0.16 | 0.17 | 0.38 |
| Intergenic | 0.32 | 0.18 | 0.18 | 0.32 |

From these per-codon coefficients we derived two complementary per-gene summaries that capture different facets of the same underlying selection regime: a translational load measure, *L*_*ROC*_, and a sign-aware efficiency measure, *ROC*_*eff*_. The two are equally well-defined statistics of the ROC posterior, but they answer different questions, and only *S*_*η*_ is a per-gene mirror of the classical Wright (1937) selection coefficient.

Translational Load (*L*_*ROC*_). A φ-weighted, magnitude-only summary of the cumulative cost of suboptimal codon choice within a gene:

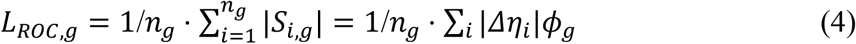

Codon-usage efficacy (*ROC*_*eff*_). A sign-aware summary of how strongly a gene’s 4-fold-degenerate sites use the preferred codon set, computed as the negative η-weighted codon usage at strictly 4-fold positions:

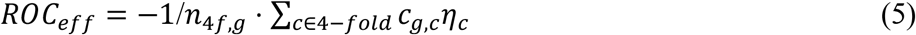

Where *c*_*g,c*_ is the count of codon c in gene g and the total number of 4-fold codons. Preferred codons have *η*_*c*_ < 0, so *ROC*_*eff*_ is positive for genes that use them preferentially and negative for genes biased toward unpreferred codons. *ROC*_*eff*_ does not involve φ (it isolates the codon-usage signal from expression-driven amplification and crucially preserves direction). At the gene level it is the closest ROC-derived counterpart to Wright’s S in the two-allele MSD framework introduced below (Spearman ρ_S ≈ +0.78; Figure S9). This metric purpose is to validate the ROC results by relating it directly to the population level selection coefficient derived from Wright 2-allele system of equations.

### Inference of optimal codons

We identified preferred codons using our best ROC-SEMPPR models, selecting the codon with the lowest estimated cost. This cost-based inference has the advantage of not requiring the assumption that selection magnitude is uniform across genes (Gilchrist et al., 2015). Unlike the standard “optimal codon” approach—which simply picks the most frequent codon among the top 50–100 most highly expressed genes—the ROC-SEMPPR framework explicitly models the opposing forces of mutation and selection. This matters because, in a genome with strong AT-mutational bias like *M. guttatus*, the most frequent codon at synonymous sites in highly expressed genes may still be mutation-driven (A/T-ending) rather than selection-driven. The ROC model disentangles these forces, identifying the codon with the lowest translational cost irrespective of its absolute frequency, and thus avoids the bias inherent in frequency-based heuristics.

### Relationship between tRNA abundance and codon/amino acid usage

To investigate the co-adaptation between synonymous codon usage and the translational machinery, we analyzed the correlation between codon frequencies and predicted tRNA availability within amino acids using Spearman’s rank correlation tests. tRNA availability was quantified using genomic gene copy number, extracted from the *Mimulus guttatus* v2.1 genome using tRNAscan-SE (Chan et al., 2021). The use of copy tRNA copy number as proxy of relative abundances of the molecular species have been validated empirically before (Camiolo et al., 2012). To distinguish between background mutation biases and selection for translational efficiency, these correlations were performed on the entire gene set as well as a subset restricted to the top 5% of most highly expressed genes, where translational constraints are expected to be strongest. As a validation of global proteomic adaptation, we further assessed the relationship between the total frequency of each amino acid in the proteome and the aggregate genomic copy number of its corresponding isoaccepting tRNAs.

To distinguish between selection for translational speed and accuracy, we examined the pairing mechanics of the optimal codons identified by the ROC-SEMPPR model. Preferred codons—those with the lowest estimated translational inefficiency (Δ*η*)— were compared to the anticodons of the most abundant tRNA species for each amino acid. We classified these interactions as either strict Watson-Crick pairings or wobble pairings (Murphy & Ramakrishnan, 2004). According to the translational accuracy hypothesis, optimal codons should preferentially form high-fidelity Watson-Crick interactions at the third codon position.

### Polymorphism analysis

To isolate the relative contributions of mutational bias and natural selection to nucleotide diversity (*π*), we partitioned the genome into distinct functional compartments: intergenic regions (analyzed globally, as well as specifically within 10 kb and 20 kb of transcription start sites), introns, and coding exons. The latter was further subdivided by positional hierarchy (first and second codon positions versus third) and site degeneracy at the third position. Nucleotide diversity (*π*) was calculated within all compartments at a site-specific level, which was essentially for comparing *π* values between compartments (e.g. 4-fold-degenerate *π* versus non-synonymous *π*), among genes within compartments (e.g. 4-fold *π* across genes that vary in expression), and among codons that vary in position (e.g. 4-fold *π* as a function of distance from gene start site).

We predicted the gene-specific frequencies of each nucleotide at 4-fold degenerate sites from gene expression levels using Generalized Additive Models (GAMs) with a Beta error distribution and logit link function. The spatial analysis was restricted to the first 200 codons to capture initiation dynamics. Additionally, to evaluate the interacting effects of gene length and expression level on the efficacy of selection, we modeled their joint influence on the global frequency of preferred codons. This was achieved by fitting 2D tensor product smooths within the GAM framework, allowing us to capture complex, non-linear interactions between log-transformed expression and length.

### Estimation from Wright two-allele mutation–selection–drift equation

The ROC-SEMPPR model described above estimates per-gene selection parameters from observed CUB combined with gene expression data. We obtained a distinct set of estimates using polymorphism data. Wright (1931) derived the density function (*ϕ*) for the allele frequency (*p*) of a favorable allele in a two-allele model under the joint action of mutation, selection and genetic drift. This can be applied to 4-fold degenerate codon sites by binning the three unpreferred codons into a single detrimental allele (McVean & Charlesworth 1999). We estimate the mutational parameters of Wright’s equation (below) from polymorphism and base frequency data within putatively neutral intron sequences. We then infer gene specific selection from the observed frequency of preferred bases at 4-fold sites within exons of that gene. The density function for p is a function of three scaled constants:

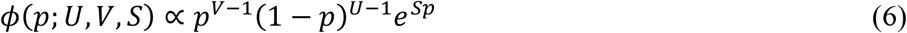

with *U* = 4*N*_*e*_ *μ, V* = 4*N*_*e*_ *ν* and *S* = 4*N*_*e*_ *s*, where μ is the mutation rate (preferred → unpreferred) and ν is the reverse rate (unpreferred → preferred). When S = 0, eq (6) becomes a beta density function and the expected values for p and 2p(1-p) have simple solutions:

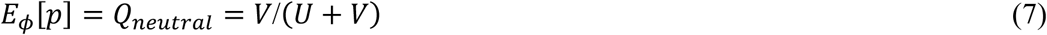

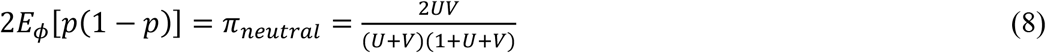

To estimate U and V, we use the full site frequency spectrum at intronic positions across the 187 sequenced IM lines. S(n, k) is the number of intronic sites where the favored base is carried by k of n lines. For estimation, we apply the Beta–Binomial likelihood (Hartl et al., 1994; Vogl, 2014):

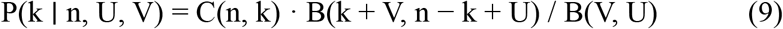

and obtain U and V by maximizing the weighted log-likelihood:

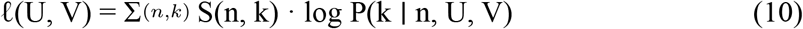

We use Nelder–Mead optimization in log(U), log(V) through the stats::optim() function in R. We use C as the “preferred” base within introns because 7 of 8 4-fold amino acids has a C-ending preferred codon. The fit yielded *V*_*C*_ = 0.00509 and *U*_*C*_ = 0.02873. Using these values in eqs (7) and (8) predict values that are close to the observed statistics.

When S ≠ 0, the equilibrium preferred-base frequency Q(S;U,V) and the expected nucleotide diversity π(S;U,V) admit closed forms of Kummer’s confluent hypergeometric function _1_*F*_1_(⋅) which we calculated in R using gsl::hyperg_1F1. For each gene g, we inverted the observed preferred-base frequency (*Q*_*g*_ computed from all 4-fold degenerate sites) on the closed-form Q(S;U,V) curve via signed bracket search:

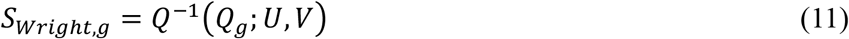

retaining sign so that *S*_*Wright,g*_ < 0 flags genes whose 4-fold pool fall below the neutral preferred-base frequency *Q*_*neutral*_ = *V*/(*U* + *V*) — i.e., genes where sampling error dominates.

## RESULTS

### CUB metrics indicate strong codon usage bias related to gene expression

Genome-wide, there is strong signature of non-random usage of synonymous codons (*G*_*genome*‐*wide*_ = 331589.8, *df* = 38, *p* < 2.2 · 10^−16^). The Effective Number of Codons (ENC) ranged between 31.8 and 59.0 (mean = 53.4), and gene-to-gene, 80.2% of genes (20213 out of 25188) show evidence of deviations from equal usage (*p* < 0.05). G tests were highly significant (207.2 ≤ *G*_*aa*_ ≤ 70177.8) for all 18 amino acids coded by at least two codons(in all cases). While genome-wide RSCU values for certain amino acids, such as Phenylalanine (Phe) and Histidine (His), appear to approach parity (Fig. 1A), there is significant intragenomic heterogeneity. The results of the G-test for heterogeneity (Table S1) demonstrate that synonymous codon usage is significantly non-uniform across the *M. guttatus* transcriptome (*p* < 0.001 for all amino acids), suggesting that individual genes are subject to varying intensities of selective and mutational pressure. CDC ranged from 0.0401 to 0.5427, with a mean/median value of 0.1266/0.1143. According to this metric 18,722 (FDR < 0.05) out of 22,556 genes have some degree of CUB (∼83%), consistent with results from the other metrics. Additionally, Fig. S2 also shows that most of the significant genes according to CDC are under the curve in the ENC plot (*χ*^2^ = 391.19, *p* = 4.56 ⋅ 10^−87^).

**Figure 1:**
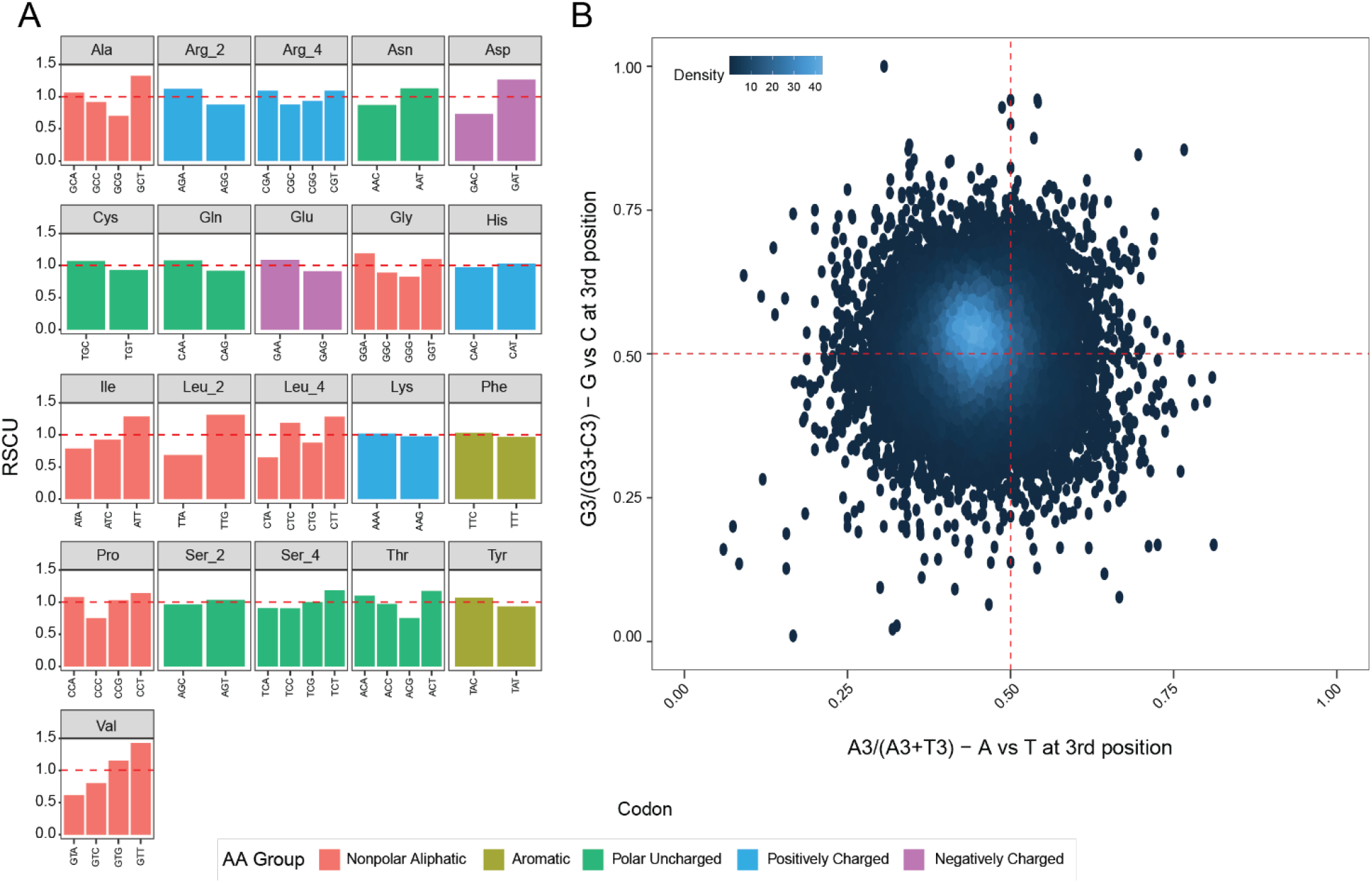
A) Relative Synonymous Codon Usage (RSCU) is the observed frequency of codons (within an amino acid) relative to equal usage. Each panel represents an amino acid set of codons, except for 6-fold amino acids, that are split in the corresponding 2- and 4-fold groups (Arg, Leu, and Ser). B) The “Parity rule” plot, with each gene as a point, depicts strand specific bias in terms of nucleotide composition. The dense core is displaced toward lower x-axis and higher y-axis values indicates a substantial T-over-A bias and more modest G-over-C bias.

To evaluate the extent of codon bias relative to genomewide base composition, we analyzed the relationship between ENC and the GC content at synonymous third positions (*GC*_3*s*_) (Fig S1B). We compared the observed data the theoretical expectation (F. Wright, 1990) if synonymous codon usage were governed solely by *GC*_3*s*_ under the assumption of parity (i.e., *A* = *T* and *G* = *C*):

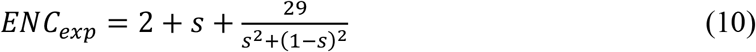

where *s* represents the observed *GC*_3*s*_ fraction. We observed that most *M. guttatus* genes fall below expectation.

Within introns, there is a notable strand bias in base frequencies: T > A > G > C (Table 1). As expected, there is no strand-specific imbalance in intergenic regions where nucleotide composition adheres to the parity rule. The observed T-bias in introns is consistent with the pattern seen in Fig. 1B. This suggests that the true mutation-drift equilibrium, which is our null model in testing for selection, is distinct from this theoretical prediction (eq 10). The ROC-SEMPPR modeling (described below) that we use to estimate codon usage selection does not assume mutational parity.

We further investigated strand-specific mutational pressures using a “Parity Rule 2” plot (Fig. 1B), which evaluates the proportionality of synonymous nucleotides within the same DNA strand (Sueoka, 1995). We plotted the intra-strand A/T ratio (*A*_3_/[*A*_3_ + *T*_3_]) against the G/C ratio (*G*_3_/[*G*_3_ + *C*_3_]) for the third codon position. With intra-strand parity—where mutation and repair rates are perfectly balanced between strands—genes are expected to cluster at the coordinates (0.5, 0.5). Contrary to this expectation, we see significant strand bias, particularly for A < T (x-axis in Fig 1B). This asymmetry suggests a strand-specific mutational bias, as might be caused by asymmetric deamination or replication-associated repair, favoring T-ending codons (U-ending in the mRNA), in agreement with Table 1.

Next, we examined the relationship between CUB metrics and gene expression (Fig. 2). Per gene values for CAI ranged from 0.54 to 0.89 (mean = 0.70) and were significantly elevated in highly expressed genes (top 5%; Fig. S3). We found that the interaction of maximum expression and expression breadth, controlling for gene length significantly predicts CDC. The GAM model explained 54% of the deviance, with both the interaction term and gene length highly significant (*p* < 2.2 ⋅ 10^−16^). CDC increases with maximum gene expression (across tissues), although the strength of the increase changes with breadth (Fig. 2). Additionally, after binning genes in three categories (top 5%, middle 90%, and bottom 5%) we saw a significant differentiation (*Kruskal−Wallis* = 196.191, *df* = 2, p < 2.2-10^− 16^; Fig. S7A).

**Figure 2:**
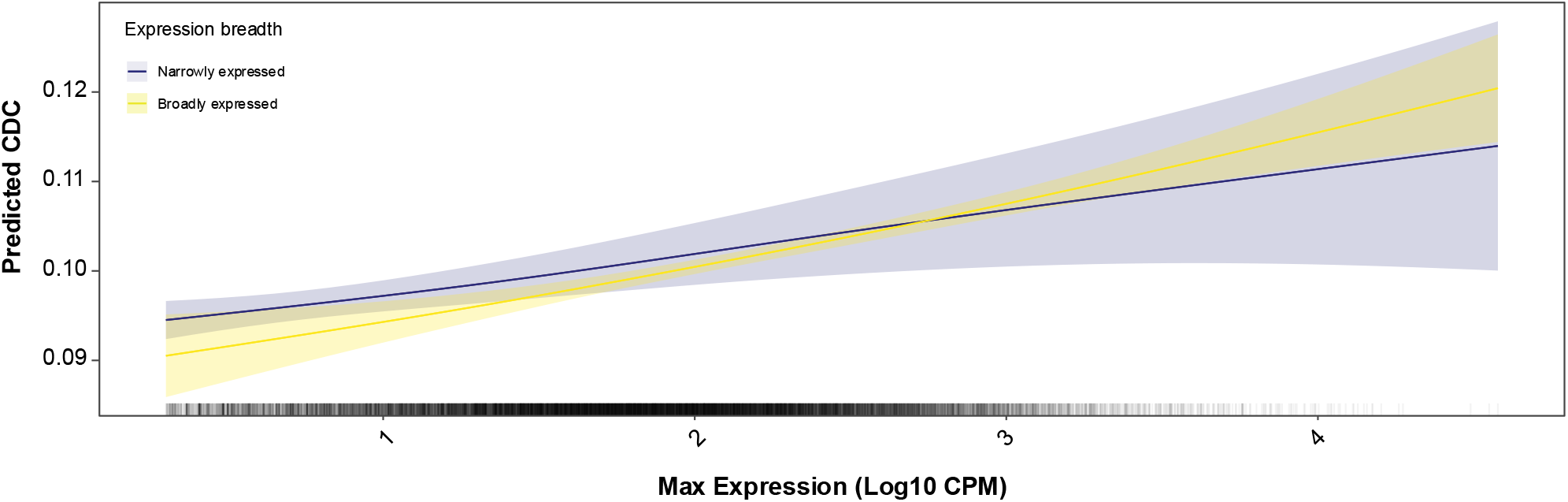
The Codon Deviation Coefficient (CDC) increases with gene expression level, and the relationship is stronger for broadly expressed than narrowly expressed genes. The lines are the GAM prediction for CDC as a function of maximum expression (log 10) and the expression breadth, controlling for gene length. The dashes over the x axis: maximum expression for each gene. Expression breadth refers to the number of tissues where CPM was higher than 1. For simplicity, we shoed the curves for genes were expression breadth was 1 (blue) and 29 (yellow).

The multivariate ordinations based on correspondence and principal component analyses place each gene as a point in a space defined by its codon usage profile; genes with similar usage cluster together. We found differences between expression groups in the multivariate spaces defined by the three main axis from correspondence analysis (*Pillai* = 0.05, *F*_(3,1915)_ = 33.78, *p* < 2.2 ⋅ 10^−16^) (Fig. S8A), and the principal components (*Pillai* = 0.20, *F*_(3,1915)_ = 163.04, *p* < 2.2 ⋅ 10^−16^) (Fig. S8B). In both cases we visualized a directional trend regarding the codons that are preferred, superimposed to the general AT-GC intragenomic bias at synonymous positions.

### ROC-SEMPPR modeling identifies preferred codons experiencing CUS in highly expressed genes

All codons favored by CUS are GC-ending and 14 out of 19 are C-ending (the list is reported in Fig. 3, alongside the preferred codons in other model plants). Figure 4 illustrates representative examples of how the ROC-SEMPPR model effectively recapitulates the transition in codon usage as a function of estimated protein synthesis rates (which is determined by gene expression levels). With low expression, predicted frequencies closely align with the mutational baseline (*ΔM*), whereas in highly expressed genes, frequencies converge toward the selective optimum (*Δη*). When mutation and selection oppose each other (mutation favors T-ending codons while selection favors C-ending codons) the predicted trajectory for the preferred codon rises from a low value at the left (mutation-dominated regime) to a high value at the right (selection-dominated regime). This produces the characteristic sigmoidal pattern for most amino acids. Apparent local inversions between predicted trajectories and observed points (most notable in Cysteine) reflect the constraint that ΔM is fixed to intronic values and cannot be adjusted to match within-coding sequence data.

**Figure 3:**
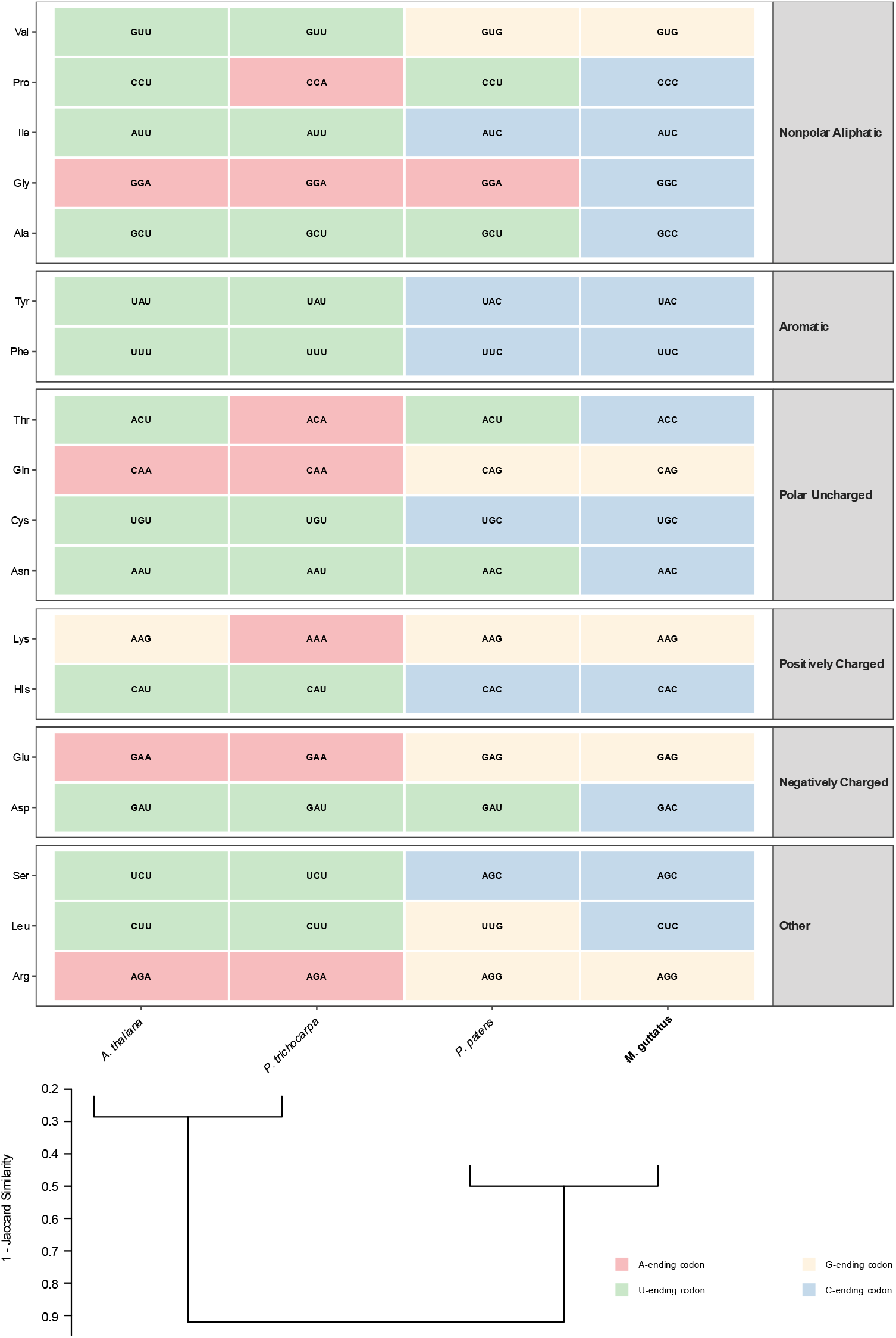
Preferred codons of *Mimulus guttatus* are compared to other plant species. Amino acid chemical family is shown to the rightmost. Colors distinguish the third nucleotide of codons.

**Figure 4:**
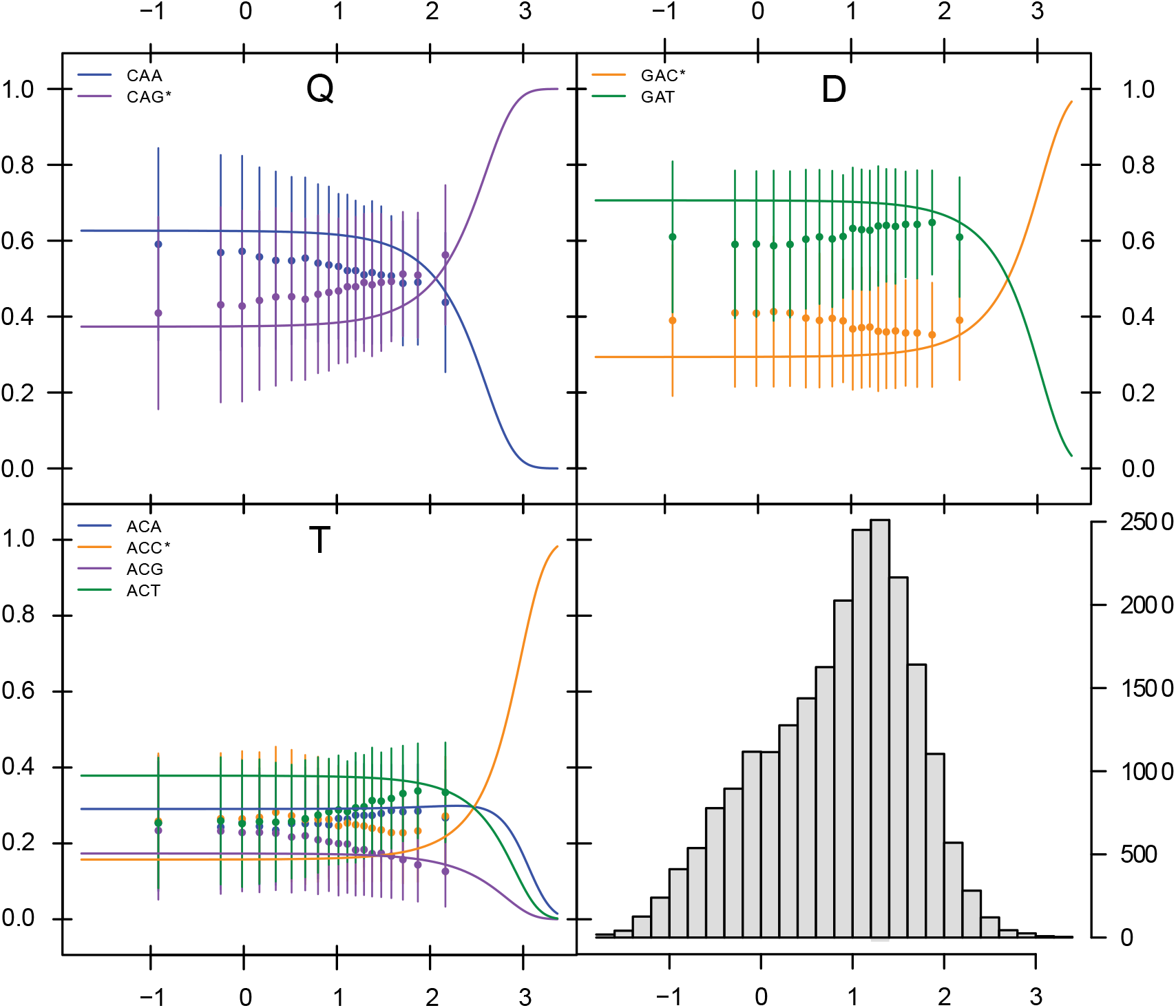
Codon frequency dynamics as a function of estimated protein synthesis rate (φ) for three of the 19 multi-codon amino acids in *M. guttatus*. The first three panels illustrate one amino acid. Points and vertical bars represent the observed mean codon frequency (±95% CI) for genes binned by their estimated protein synthesis rate φ. Solid colored lines show the predicted codon frequency trajectories from the best-fitting ROC-SEMPPR model (dM-fixed-with-φ-introns), in which mutational bias (ΔM) is fixed from intronic nucleotide frequencies and protein synthesis rates are informed by empirical expression data. The histogram (lower right panel) shows the distribution of φ values across the genome.

While the model demonstrates high overall performance (based on convergence and stability of the log likelihood and posterior traces for the estimated parameters), we observed localized deviations where predicted frequencies do not mirror observations (e.g., the subtle inversion in cysteine codons at low and middle expression). These mismatches are biologically instructive and likely reflect the discrepancy between contemporary expression snapshots and the long-term evolutionary pressures that have shaped the genome. Furthermore, such instances that escape the multinomial prediction could be due to the noisiness associated with expression data or could even suggest that additional layers of selection—such as constraints on mRNA secondary structure or localized tRNA pool fluctuations—may be operating alongside translational efficiency to fine-tune the *Mimulus* coding landscape. Overall however, there is a negative correlation between ROC cost parameters (Δη) and expression-driven frequency changes across all 59 codons (ρ = −0.329, p = 0.011), which confirms that the ROC model is capturing genuine selective signal.

### Wright’s equation suggests that CUS affects many genes, but that selection is usually weak

Polarizing 4-fold sites based the preferred codon inferences from the ROC-SEMPPR model, selection coefficients inferred from Wright’s equation, *S*_*Wright*_, ranged between −2.01 and 2.84 (Mean and median were 0.61 and 0.56 respectively Figure 5A). *S*_*Wright*_ estimates from a gene are inferred based on the observed frequency of preferred codons combined with the mutation rates estimates from introns (U = 0.0297, V = 0.0051). As expected, U and V imply a strong mutational bias away from the preferred base. 4,029 genes have point estimates for *S*_*Wright*_ ≥ 1, 16,179 genes have *S*_*Wright*_ between 0 and 1, and 1,133 genes have negative values. We roughly categorized these as “drift dominated” (*S*_*Wright*_ ≤ 0), “weakly selected*”* (0 < *S*_*Wright*_ < 1), and “strongly selected” (*S*_*Wright*_ ≥ 1). Importantly, there must be considerable estimation error in gene-specific *S*_*Wright*_ owing to the limited number of 4-fold sites per gene (168 on average). Simple binomial sampling on the number of 4-fold sites that are polymorphic within gene (around the expected value given the true *S*_*Wright*_) will generate non-trivial error in the estimated *S*_*Wright*_. Even after allowing for this variation however, it is clear that a substantial fraction of genes are affected by CUS, particularly in the range of 0 < *S*_*Wright*_ < 1.

**Figure 5:**
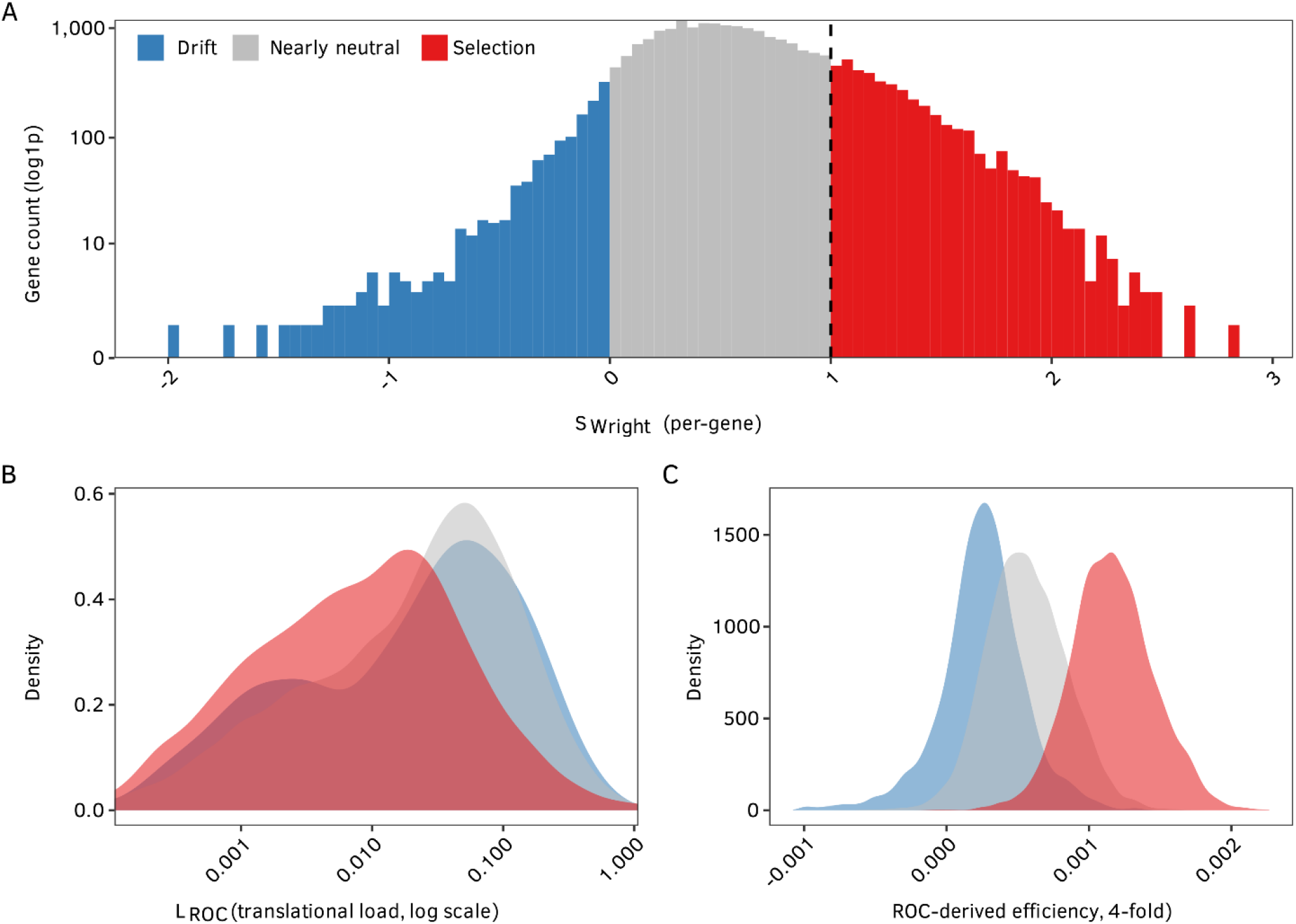
A) Distribution of *S*_*Wright*_ values across genes based on polymorphism data from Iron Mountain, Oregon. The dashed line is the drift barrier threshold (S = 1). Lower panels: Distributions of *L*_*ROC*_ (B) and *ROC*_*eff*_ (C) across genes that were classified as drift (blue), nearly neutral (grey), or selection (red) using Wright’s formula.

There is a compelling correspondence between *S*_*Wright*_and the ROC-SEMPPR estimates obtained from the same genes (Fig 5B,C). Genes that ROC-SEMPPR predicts to have lower translational load are much more likely to have *S*_*Wright*_ ≥ 1 (Fig. 5B). The ROC-SEMPPR classification of efficiency at 4-fold codons (in terms of gene specific usage) increases continuously from drift dominated to Nearly neutral to Selection dominated according to Wright-based estimates (Fig. 5C). We found a highly significant enrichment for specific GO terms in genes where *S*_*Wright*_ > 1 (Table S2). In most cases, those terms were associated with the molecular machinery of photosynthesis (*e*.*g*. photosystem, thylakoid), translation, phosphate scavenging, defense mechanisms, and cell wall biogenesis and remodeling. Among the genes with the highest estimated *L*_*ROC*_ are Light-Harvesting Complex II (MgIM767.03G128800, LHCII 5) and Ribulose bisphosphate carboxylase small chain (MgIM767.09G049300). Table S3 provides a complete list matched to homologs in *Arabidopsis thaliana*.

### Preferred codon frequency shows a strong dependency on position within a gene

There is a highly significant, but non-monotonic, relationship between preferred codon frequency and distance from the gene start site (Fig. 4A). For all expression levels, the peak occurs around the 50^th^ codon and decays after that. As described below in the Discussion, we hypothesize this non-monotonic pattern results from combined effect the “translational ramp” effect (Kozak, 1999; Tuller et al., 2010) on CUS combined with a pervasive effect of linked selection. Additionally, the frequencies of the preferred codons increase with gene expression level for both broadly and narrowly expressed genes. Figure 6B illustrates GAM model predictions for preferred codon frequency controlling as a function of maximum expression, the expression breadth, and the gene length. Highly expressed genes have more preferred codons than the rest of the genes in the genome (Fig. S7B).

**Figure 6:**
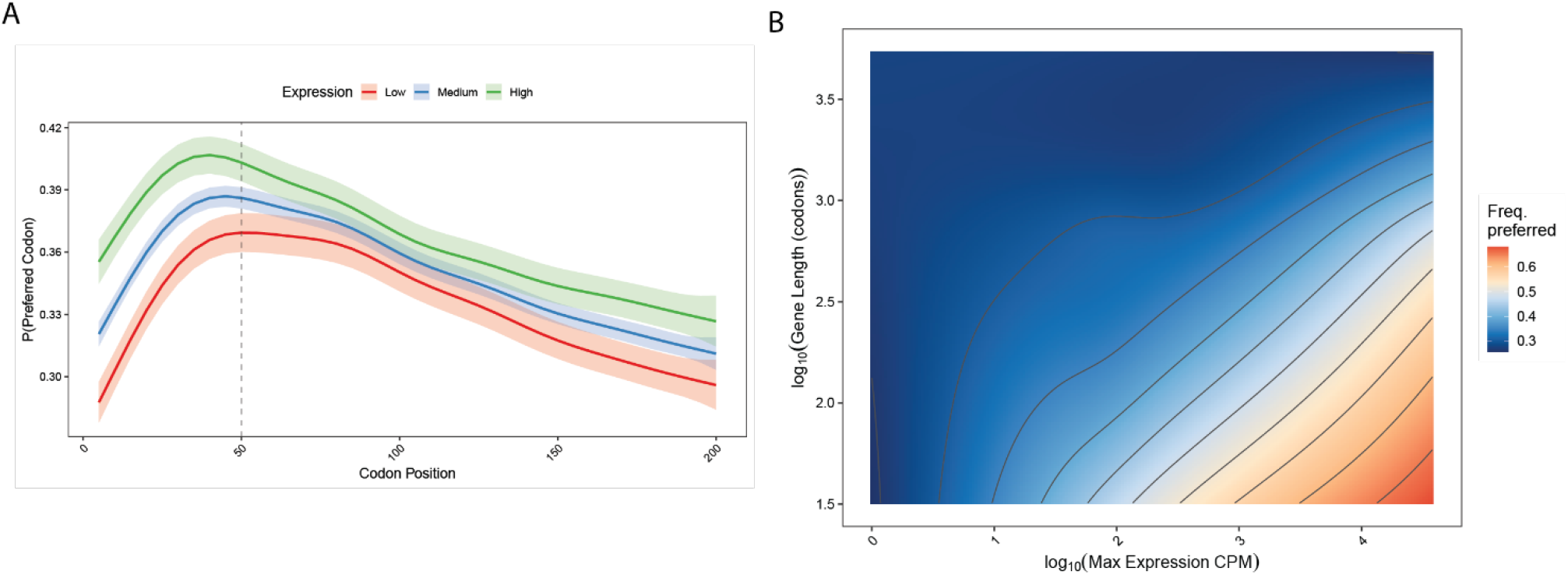
Preferred codon frequency as a function of (A) codon position across the first 200 codons of each gene. For (A), there are three different trajectories, which correspond to the low expression group (Bottom 5%), the medium expression group (Middle 90%), and the high expression group (Top 5%). B) Landscape of preferred codon frequency (4-fold sites) as a function of gene length and maximum expression.

We found no significant association between the identity of the ROC-preferred G- or C- ending codon and Watson-Crick (versus wobble) pairing at the third codon position (Fishers exact *p* = 0.17). This does not support the translation accuracy hypothesis (Akashi, 1994; Chan et al., 2017; M. Sun & Zhang, 2022). We also examined the correspondence between codon usage and the tRNA gene pool (Fig. S5A). Finally, we observed a powerful correlation between total amino acid frequencies derived from the reference transcriptome and tRNA supply (r = 0.761), indicating that the overall *M. guttatus* translational machinery is scaled to match broad proteomic demand (Fig. S5B). However, when examining individual codon-tRNA dynamics, tRNA gene copy numbers for the optimal C- and G-ending codons are generally comparable to, and in some cases lower than, those for A- and T-ending codon (non-statistically significant test).

### Synonymous polymorphism exhibits an intragenic signal of linked selection and a diversity enhancing effect of CUS

Nucleotide diversity *usually* declines as gene expression increases at both synonymous and non-synonymous sites (Fig 7A, note difference in scale given that mean 4-fold diversity is 5 times higher than non-synonymous diversity). However, there is a remarkable reversal in the trend for synonymous variation which increases in highly expressed genes (Fig. 7A), which parallels a similar pattern is the average frequency of the preferred codon per gene (Fig. 7B). As expected, nucleotide diversity is elevated near the start of genes (where recombination is high) relative to the middle of gene bodies (Fig 7C). At a given physical distance from the gene start, diversity is higher at synonymous sites than within introns, although the difference declines as we move away from the gene start. Given that introns are our neutral proxy, the analysis indicates that CUS is generally increasing nucleotide diversity at synonymous sites (Fig. 7D). Given the strong mutational bias towards unpreferred bases at 4-fold sites, diversity is predicted to increase over the range from S > 0 to S = 1.5 (McVean & Charlesworth, 1999).

**Figure 7:**
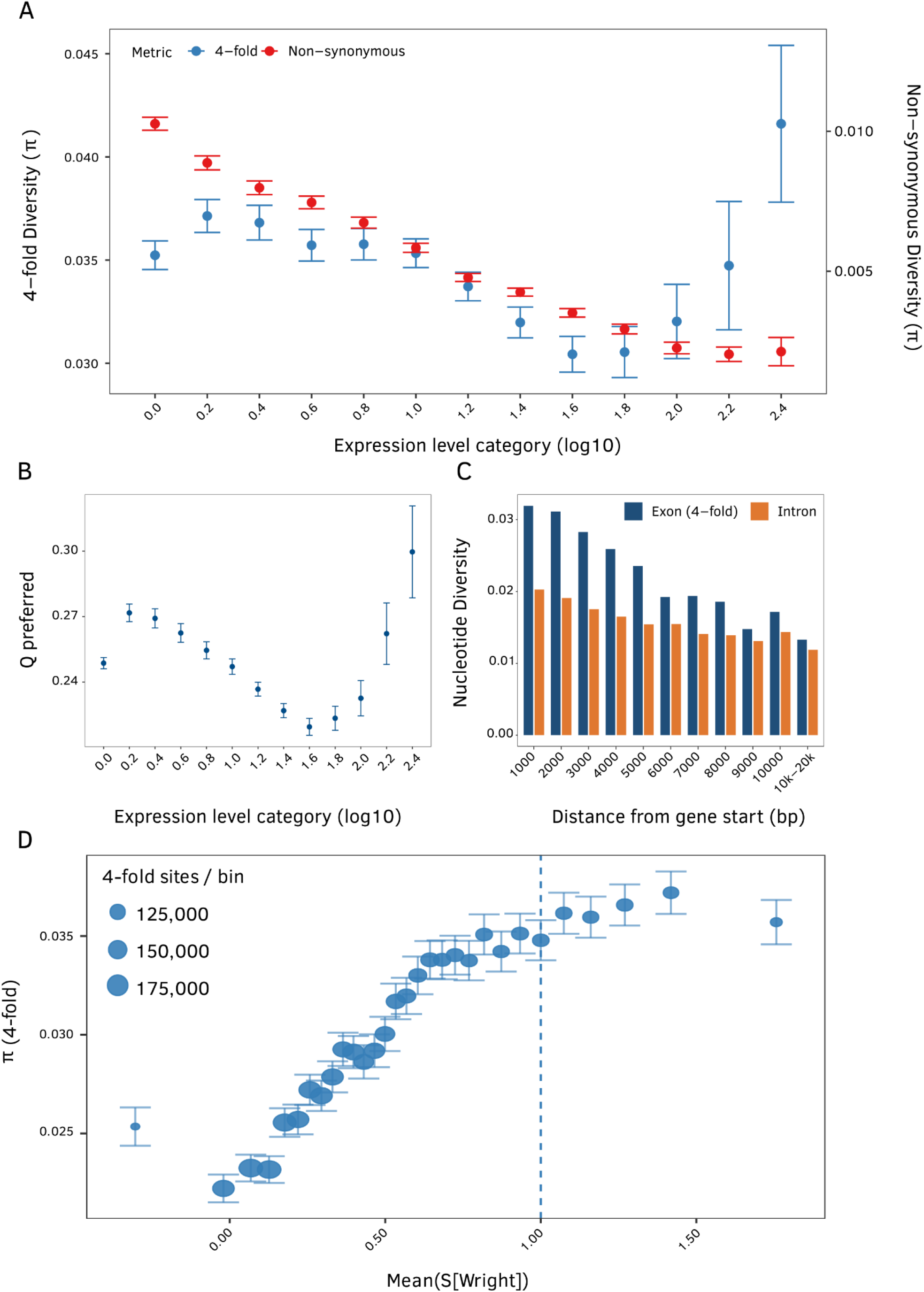
Patterns of nucleotide diversity associated with codon usage selection (CUS). A) Non-synonymous and synonymous nucleotide diversity as a function of expression level. B) Frequency of the preferred codon as a function of expression levels. Nucleotide diversity of introns and exons (synonymous sites) across gene bodies. D) Synonymous nucleotide diversity as a function of the population level selection coefficient derived from the two-allele model of Wright.

## DISCUSSION

Gene expression level is an important determinant of nucleotide variation in genes of *M. guttatus* via two important but opposing mechanisms. Purifying selection on amino acid changes appears to increase with gene expression level which depresses linked synonymous variation through background selection (Charlesworth et al., 1993). However, increasing gene expression also strengthens codon usage selection (CUS). For highly expressed genes, such as the main constituents of the photosynthetic machinery, CUS is strong enough to counteract background selection, increasing synonymous variation even when nonsynonymous variation reaches its minimum (Fig 7A). Below, we first review the population genetic aspects of simultaneous CUS and linked selection and then discuss the mechanistic aspects of codon usage in the genes most affected by translational selection.

### Population-genetic signatures of background selection and codon selection

Increasing purifying selection with higher gene expression has been demonstrated in numerous species based on both inter- and intra-specific data. There is a very general observation that nonsynonymous sequence divergence between closely related species is negatively correlated with gene expression level (Pál et al., 2001; J. Zhang & Yang, 2015). We find a striking negative relationship between non-synonymous π and the expression level of genes (Fig 7A) suggesting that purifying selection on amino acid changes is much stronger in highly expressed genes (Trucchi et al., 2023). Synonymous π at 4-fold degenerate sites follows a parallel decline over most of the expression range. This general decline in both site classes is most parsimoniously explained by background selection (Charlesworth et al., 1993)—purifying selection on nonsynonymous mutations reduces variation at closely linked synonymous sites. Local recombination rate is critical here. In *M. guttatus* and numerous other plants, recombination rate is high near gene start sites and declines toward the middle of genes (Hellsten, et al., 2013). This recombination gradient should create a spatial gradient in the efficacy of selection within genes. Fig 7C provides direct evidence for linked selection: Synonymous variation is highest where recombination is highest and declines as recombination decreases in frequency.

The population genetic signal of CUS, where synonymous polymorphisms are directly selected is subtle but multi-faceted. If synonymous variation were entirely neutral, both synonymous and nonsynonymous π would decline as expression increases (as background selection increases). Instead, synonymous π shows a clear upward inflection at the highest expression levels (Fig 7A), which is mirrored by an increase in the average frequency of the preferred codon per gene (Fig 7B). Estimates for the strength of CUS suggest that most genes are in the range of 0 < S < 1 (Fig 5A). 82% of genes (18,273 of 22,355) fall below the so-called drift threshold (*S* < 1), suggesting a dominant effect of mutation and drift. While *S* < 1 is often classified as ‘nearly neutral’, CUS can have an important quantitative effects over this range. Given our estimates for U and V, the predicted frequency of the preferred codon doubles between S = 0 and S = 1. Importantly, over the entire range from S=0 to S=1, synonymous diversity is predicted to increase. McVean & Charlesworth (1999) showed that when selection favors one allele, but the opposing mutational pressure is sufficiently strong, diversity will be greatest at intermediate values of S. When mutation and selection are evenly balanced, the result is an excess of heterozygosity relative to neutral sites. Fig 7D suggests that much of the *M. guttatus* genome is on the upslope of this diversity hump.

In several important ways, we may be underestimating CUS. Most basically, our fitting of Wright’s equation to observed preferred codon frequencies per gene assumes there is no effect of background selection or Hill-Robertson interference affecting synonymous variants (Hill & Robertson, 1966). In so far as these processes are important, *S*_*Wright*_will underestimate S. Also, our modeling has assumed that a single nucleotide base is preferred per codon, regardless of the gene or position within gene. The “translational ramp” hypothesizes an advantage to placing sub-optimal codons at the 5’ end of genes (just following the start codon) to slow early elongation in translation. This allows proper spacing of trailing ribosomes to prevent downstream collisions and give the nascent peptide time to engage the signal recognition particle for membrane targeting (Kozak, 1999; Tuller et al., 2010). These principles are now routinely applied in genetic engineering to enhance the expression of heterologous genes (Parvathy et al., 2022). “Sub-optimal” in this context refers only to elongation speed and not overall fitness.

Fig 6A is fully consistent with the translational ramp hypothesis. Just after the start codon is where we expect the highest frequency of favored codons if considering only the effect of Hill-Robertson interference or background selection (that is where recombination rate is highest). However, preferred base frequency increases over the first 50 codons across all expression categories of genes. From a population genetic perspective, the implication is that we are failing to recognize that selection is heterogeneous across sites. Fitting a model with a single constant S per codon, and treating all alternative codons as equally determinantal will underestimate the true strength of selection.

A second important assumption of our analysis is that we assumed mutation to be homogeneous within and across genes – our application of Wright’s equation used a single U (0.0287) and V (0.0051). Among other mutational processes, the frequency of GC-biased gene conversion could vary over the genome. During meiotic recombination, heteroduplex repair can systematically favor G/C over A/T alleles, which is the opposite of the overall mutational bias towards A/T. Our main evidence for CUS is how the frequency of C changes at degenerate positions as a function of gene expression level (Fig 7B). We have no evidence that the rate gene conversion changes within gene expression in the same way, but more work is needed on the fine scale localization of both crossover and gene conversion events in *M. guttatus*.

### Codon selection is driven genes in regulation, translation, structure, and photosynthesis

Standard heuristic codon usage indices (CAI, CDC) show significant but weak correlations with gene-specific *S*_*Wright*_ and *ROC*_*eff*_ estimates. This indicates that metrics which conflate mutational bias with translational selection, or which fail to account for the directionality of fitness, are insufficient for quantifying translational selection in *M. guttatus*. The ROC-SEMPPR and *S*_*Wright*_ analyses deconvolve these evolutionary forces, placing the results from *M. guttatus* in an informative comparative context. In *E. coli* and *S. cerevisiae*, translational selection shapes codon usage across a substantial fraction of the genome, and preferred codons closely match the most abundant tRNAs (Gouy & Gautier, 1982; Akashi, 1994; Gilchrist et al., 2015). In *Drosophila melanogaster*, CUS is evident for many genes, though weaker than in unicellular organisms, with selection coefficients estimated to be on the order of for a meaningful fraction of the transcriptome (Machado et al., 2020). At the other end of the spectrum, codon usage in rice and maize appears to be driven primarily by mutational bias and genome-wide GC content variation, with only weak evidence for translational selection (Wang & Hickey, 2007; Campbell & Gowri, 1990). *M. guttatus*, with its ∼19% of genes exceeding the drift threshold (4*N*_*e*_ *s* > 1), occupies an intermediate position: translational selection is clearly operating but is confined to a modest subset of the genome.

Previous CUB research in plants has focused on two species from the Rosid clade (*A. thaliana* and *P. trichocarpa*), the moss *P. patens* (Parvathy et al., 2022), and the two primary monocot crops, corn and rice. Our analysis of *M. guttatus*, an Asterid, produces qualitatively different patterns. notably, the preferred codons are predominantly C-ending (74% of preferred codons; Fig. 3), rather than the A/U-ending codons favored in the Rosid species. A straightforward explanation would be that C/G-ending codons are preferred because their cognate tRNAs are more abundant. However, we found no evidence for a disproportionately large tRNA gene pool matching C- or G-ending codons (Fig. S5A) This apparent paradox dissolves when considered in the broader genomic context. Because 81% of the transcriptome is experiencing weak selection, most of its evolution is shaped by drift and mutational bias toward A/T (Table 1), the global tRNA pool has likely co-evolved to service this AT-biased demand. The overall amino acid–tRNA correspondence is strong (Spearman *r* = 0.761, *p* = 3.83 × 10^−4^; Fig. S5B), indicating that the *M. guttatus* translational machinery is broadly scaled to match proteomic demand. But within amino acid families, tRNA gene copy numbers for optimal C- and G-ending codons are generally comparable to, and sometimes lower than, those for A- and T-ending codons. Furthermore, a Fisher’s exact test found no significant association between the identity of the ROC-preferred codon and Watson-Crick (versus wobble) pairing at the third codon position (*p* = 0.17), providing no support for the translational accuracy hypothesis. This distinction underscores that the identity of preferred codons is not universally conserved among plants but reflects lineage-specific interactions between the mutational landscape, tRNA pool, and the intrinsic kinetic properties of codon–anticodon interactions.

Interpreting the GO enrichment results reveals a coherent and biologically coherent picture of the selective pressures acting on this gene set. Rather than a diffuse signal, the data converge on a set of interconnected functional priorities: the regulation of enzymatic activity, the maintenance of translational capacity, the dynamic remodeling of the cell wall, the selective trafficking of ions and metabolites, and the support of photosynthetic function. Five overlapping themes emerge from the enrichment landscape:

#### Enzyme Regulation and Inhibition

The most statistically compelling signals in the dataset concern the modulation of cellular activity rather than catalysis itself. Molecular function inhibitor activity (GO:0140678; p = 6.94×10^−18^) and enzyme inhibitor activity (GO:0004857; p = 1.15×10^−17^) rank among the top terms genome-wide, with enzyme regulator activity (GO:0030234) close behind. This pattern suggests strong selection on the fine-tuning of metabolic flux, likely through post-translational control of enzyme output rather than changes in enzyme abundance alone.

#### Translation and Ribosomal Integrity

The protein synthesis apparatus emerges as a second major axis of enrichment. Structural constituent of ribosome (GO:0003735) is highly significant (p = 3.58×10^−11^), and the broader term translation (GO:0006412) reinforces this signal. Complementary terms — including structural molecule activity (GO:0005198) and protein heterodimerization activity (GO:0046982) — point to selection on the physical assembly and stability of macromolecular complexes, not just the synthesis reaction itself.

#### Cell Wall Organization and Biochemical Modification

A substantial cluster of enriched terms reflects the dynamic construction and remodeling of the plant cell wall. Cell wall organization (GO:0071555) and its broader counterpart cell wall organization or biogenesis (GO:0071554) are both enriched, as are cell wall modification (GO:0042545), external encapsulating structure organization (GO:0045229), pectinesterase activity (GO:0030599), and a suite of transferase activities consistent with the enzymatic remodeling of polysaccharide and lipid components of the wall matrix.

#### Ion and Metabolite Transport

The controlled movement of small molecules across cellular and organellar membranes constitutes a distinct and broad enrichment theme. Phosphate ion transport (GO:0006817) and its transporter counterpart (GO:0005315) are among the more significant terms, accompanied by inorganic anion transport (GO:0015698) and inorganic ion transmembrane transport (GO:0098660). Nitrogen acquisition is represented by ammonium channel activity (GO:0008519) and ammonium transmembrane transport (GO:0072488), while sodium ion transport (GO:0006814) adds further breadth. These signals are complemented by secretion (GO:0046903), secretion by cell (GO:0032940), and export from cell (GO:0140352), collectively pointing to a heavily selected interface between the cell and its environment.

#### Photosynthetic Machinery, Pigments, and Redox Chemistry

The photoautotrophic functions of the cell are well-represented across multiple levels of biological organization. The overarching term photosynthesis (GO:0015979) is enriched alongside specific structural components — photosystem I (GO:0009522), photosystem I reaction center (GO:0009538), the thylakoid (GO:0009579), and the photosynthetic membrane (GO:0034357). Pigment biosynthetic process (GO:0046148) and pigment metabolic process (GO:0042440) reflect selection on the tetrapyrrole and chlorophyll pathways, corroborated by tetrapyrrole binding (GO:0046906) and heme binding (GO:0020037). Redox-related terms — electron transfer activity (GO:0009055), monooxygenase activity (GO:0004497), and oxidoreductase activities acting on paired donors (GO:0016705) or sulfur group donors (GO:0016667) — further link the photosynthetic and metabolic redox networks. Catechol oxidase activity (GO:0004097) and protein-disulfide reductase activity (GO:0015035) extend this oxidative chemistry into secondary metabolism and protein homeostasis.

Taken together, the enrichment profile is consistent with pervasive selection on the efficiency and regulation of core plant cell functions: translating proteins accurately, structuring and remodeling the wall, trafficking ions with precision, and sustaining photosynthetic output — all coordinated through tight enzymatic control.

### Genes with even small inefficiencies can have a massive translational load (*L*_*ROC*_)

Genes with elevated L_ROC values are those where the cost of translational inefficiency is highest because they are both highly expressed and encode proteins where errors or slowdowns have outsized fitness consequences. The composition of this group is therefore not random; it reflects the cell’s most critical and resource-intensive investments. The single most striking feature of this list is the concentration of components tied to light capture and energy transduction. Photosystem II subunits (O-2, P-1, R), light harvesting complex proteins of both PSI and PSII, a Rubisco small subunit (two independent paralogs), rubisco activase, fructose-bisphosphate aldolase, transketolase, and magnesium chelatase (the committed step in chlorophyll biosynthesis) are all present. These are proteins that must be produced in enormous quantities to saturate the chloroplast, and mistranslated versions that fold poorly or aggregate would directly impair photosynthetic efficiency. The presence of the plasma membrane H^+^-ATPase and multiple vacuolar ATP synthase subunits extends this bioenergetic theme beyond the chloroplast.

Molecular chaperones are also highly represented. HSP70 appears three times (HSP70, HSC70-1, and an HSP70 family protein), HSP81-3 (a cytosolic HSP90) is present, chaperonin 60β is included, and CLPC1 rounds out the group. This is notable: chaperones are themselves highly expressed precisely because they must be present in excess to handle the folding burden of the entire proteome. A mistranslated chaperone cannot perform its function, making codon usage bias in these genes doubly important. Beyond photosystems, the list captures the enzymatic machinery that fixes and processes carbon: two Rubisco small subunit genes, rubisco activase, transketolase (regeneration of ribulose-5-phosphate), and fructose-bisphosphate aldolase. This is the suite of enzymes that must operate continuously and in stoichiometric balance. Elongation factor Tu (a GTP-binding protein that delivers aminoacyl-tRNAs to the ribosome) and poly(A)-binding protein 8 both appear, pointing to selection on the translational machinery itself. The ubiquitin pathway reflects the cell’s continuous investment in clearing damaged or aberrant proteins, a burden that would be exacerbated by elevated mistranslation.

## CONCLUSIONS

Codon Usage Bias (CUB) is important because the genetic code is neither universal nor static, it continues to evolve under various drivers (Koonin & Novozhilov, 2009). Codon preferences can shift, genetic codes can evolve, and non-standard amino acids can be recruited through code expansion or reprogramming. Indeed, some authors consider the pattern of preferential codon usage as “the second genetic code” (Parvathy et al., 2022). For evolutionary biology in general, CUB provides a valuable “case study” where three of the fundamental forces (mutation, selection, and genetic drift) are comparable in strength. In *M. guttatus*, most of the transcriptome (∼76%) is weakly affected by translational selection (0 < S < 1) and 19% experience strong selection (S > 1). The preferred codon is elevated relative to neutral expectations but remains infrequent owing to mutational pressures. However, even this weak selection is important in elevating nucleotide diversity at 4-fold sites), preserving optimal alleles against a mutational headwind. The striking decay of synonymous nucleotide diversity along the gene body, as recombination declines, highlights how background selection or other forms of linked selection can choke off selection of this strength. Within a set of highly expressed genes, overwhelmingly involved in photosynthetic, metabolic, and chaperone processes, selection more effectively opposes mutation and drift, enriching G/C-ending codons to maximize translational efficiency. Considering synonymous variants across the whole of the Mimulus genome, mutation and drift usually dominate, but translational selection achieves rare, localized victories in highly expressed genes.

## ACKNOWLEDGMENTS

We acknowledge the help and support of the KU EEB Genetics group which provided feedback and recommendations in the first stages of this research. We acknowledge conversations with Justin Blumienstiel that inspired this study and thank Jae Choi for recommendations to improve the narrative. We acknowledge grants IOS-2421689 and NSF-2421327 for KU award, which supported Dr. John K. Kelly and Luis J. Madrigal-Roca, respectively.

## AUTHOR CONTRIBUTIONS

LJMR and JKK designed and planned the research, conducted the analysis, and wrote the manuscript.

## DATA AVAILABILITY

Code and primary data are available at: https://github.com/luismadrigal98/CUB_Mguttatus. VCF files are available upon request given the size.

## SUPPORTING INFORMATION

Supplementary Figure 1: General patterns of Codon Usage Bias in *Mimulus guttatus* from Iron Mountain, Oregon, USA.

Supplementary Figure 2: ENC plot showing the ENC expectation as a function of GC3s content as a black line.

Supplementary Figure 3: Codon Adaptation Index (CAI) and its relationship with expression in the yellow monkey flower, *Mimulus guttatus*, from the natural population Iron Mountain, Oregon, USA.

Supplementary Figure 4: Codon inefficiencies relative to a reference (black dots) estimated from AnaCoDa models for *Mimulus guttatus* from Iron Mountain, Oregon, USA.

Supplementary Figure 5: Co-evolution of tRNA gene supply and genomic composition in *Mimulus guttatus*.

Supplementary Figure 6: Partial Effect of Expression Intensity on Preferred Codon Frequency.

Table S1: G-based heterogeneity test for CUB across amino acid families.

Supplementary Figure 7: Codon Deviation Coefficient Residuals (CDC Residuals) and its relationship with expression in the yellow monkey flower, *Mimulus guttatus*, from the natural population Iron Mountain, Oregon, USA.

Supplementary Figure 8: Highly and lowly expressed genes in a bi-dimensional space defined by dimensions 1 and 3 of a correspondence analysis (A) based on codon counts and the two first principal components from a PCA based on the RSCU scores (B) for *Mimulus guttatus* from iron Mountain, Oregon.

Supplementary Figure 9: Relationship between ROC-derived efficiencies and the predictions of Wright equation after transforming the nucleotide composition at 4-fold sites into a two-allele system (preferred/unpreferred bases).

Table S1: G-based heterogeneity test for CUB across amino acid families.

Table S2: GO enrichment results from genes where *S*_*Wright*_ > 1 in *Mimulus guttatus* from Iron Mountain, Oregon.

Table S3: Top genes from *Mimulus guttatus* as a function of the translational load (coefficient, *L*_*ROC*_) and their best hit in *Arabidopsis thaliana*.

## REFERENCES

Akashi, H. (1994). Synonymous codon usage in Drosophila melanogaster: Natural selection and translational accuracy. Genetics, 136(3), 927–935. 10.1093/genetics/136.3.927

Begun, D. J., & Aquadro, C. F. (1992). Levels of naturally occurring DNA polymorphism correlate with recombination rates in D. melanogaster. Nature, 356(6369), 519–520. 10.1038/356519a0

Bulmer, M. (1991). The selection-mutation-drift theory of synonymous codon usage. Genetics, 129(3), 897–907. 10.1093/genetics/129.3.897

Camiolo, S., Farina, L., & Porceddu, A. (2012). The relation of codon bias to tissue-specific gene expression in Arabidopsis thaliana. Genetics, 192(2), 641–649. 10.1534/genetics.112.143677

Campbell, W. H., & Gowri, G. (1990). Codon Usage in Higher Plants, Green Algae, and Cyanobacteria. Plant Physiology, 92(1), 1–11. 10.1104/pp.92.1.1

Castellano, D., Coronado-Zamora, M., Campos, J. L., Barbadilla, A., & Eyre-Walker, A. (2016). Adaptive Evolution Is Substantially Impeded by Hill–Robertson Interference in *Drosophila*. Molecular Biology and Evolution, 33(2), 442–455. 10.1093/molbev/msv236

Chan, S., Ch’ng, J.-H., Wahlgren, M., & Thutkawkorapin, J. (2017). Frequent GU wobble pairings reduce translation efficiency in Plasmodium falciparum. Scientific Reports, 7(1), 723. 10.1038/s41598-017-00801-9

Charlesworth, B., Morgan, M. T., & Charlesworth, D. (1993). The effect of deleterious mutations on neutral molecular variation. Genetics, 134(4), 1289–1303. 10.1093/genetics/134.4.1289

Comeron, J. M., Williford, A., & Kliman, R. M. (2008). The Hill–Robertson effect: Evolutionary consequences of weak selection and linkage in finite populations. Heredity, 100(1), 19–31. 10.1038/sj.hdy.6801059

Cope, A. L., & Shah, P. (2022). Intragenomic variation in non-adaptive nucleotide biases causes underestimation of selection on synonymous codon usage. PLOS Genetics, 18(6), e1010256. 10.1371/journal.pgen.1010256

Corbett-Detig, R. B., Hartl, D. L., & Sackton, T. B. (2015). Natural Selection Constrains Neutral Diversity across A Wide Range of Species. PLOS Biology, 13(4), e1002112. 10.1371/journal.pbio.1002112

Crick, F. H. C., Barnett, L., Brenner, S., & Watts-Tobin, R. J. (1961). General Nature of the Genetic Code for Proteins. Nature, 192(4809), 1227–1232. 10.1038/1921227a0

Cutter, A. D., & Payseur, B. A. (2013). Genomic signatures of selection at linked sites: Unifying the disparity among species. Nature Reviews Genetics, 14(4), 262–274. 10.1038/nrg3425

Danecek, P., Bonfield, J. K., Liddle, J., Marshall, J., Ohan, V., Pollard, M. O., Whitwham, A., Keane, T., McCarthy, S. A., Davies, R. M., & Li, H. (2021). Twelve years of SAMtools and BCFtools. GigaScience, 10(2), giab008. 10.1093/gigascience/giab008

Deng, Y., De Lima Hedayioglu, F., Kalfon, J., Chu, D., & Von Der Haar, T. (2020). Hidden patterns of codon usage bias across kingdoms. Journal of The Royal Society Interface, 17(163), 20190819. 10.1098/rsif.2019.0819

Dinno, A. (2024). Dunn.Test: Dunn’s Test of Multiple Comparisons Using Rank Sums. 10.32614/CRAN.package.dunn.test

Galtier, N., Roux, C., Rousselle, M., Romiguier, J., Figuet, E., Glémin, S., Bierne, N., & Duret, L. (2018). Codon Usage Bias in Animals: Disentangling the Effects of Natural Selection, Effective Population Size, and GC-Biased Gene Conversion. Molecular Biology and Evolution, 35(5), 1092–1103. 10.1093/molbev/msy015

Georgakopoulos-Soares, I., Koh, G., Momen, S. E., Jiricny, J., Hemberg, M., & Nik-Zainal, S. (2020). Transcription-coupled repair and mismatch repair contribute towards preserving genome integrity at mononucleotide repeat tracts. Nature Communications, 11(1), 1980. 10.1038/s41467-020-15901-w

Gilchrist, M. A. (2007). Combining Models of Protein Translation and Population Genetics to Predict Protein Production Rates from Codon Usage Patterns. Molecular Biology and Evolution, 24(11), 2362–2372. 10.1093/molbev/msm169

Gilchrist, M. A., Chen, W.-C., Shah, P., Landerer, C. L., & Zaretzki, R. (2015). Estimating Gene Expression and Codon-Specific Translational Efficiencies, Mutation Biases, and Selection Coefficients from Genomic Data Alone. Genome Biology and Evolution, 7(6), 1559–1579. 10.1093/gbe/evv087

Gouy, M., & Gautier, C. (1982). Codon usage in bacteria: Correlation with gene expressivity. Nucleic Acids Research, 10(22), 7055–7074. 10.1093/nar/10.22.7055

Haasl, R. J., & Payseur, B. A. (2016). Fifteen years of genomewide scans for selection: Trends, lessons and unaddressed genetic sources of complication. Molecular Ecology, 25(1), 5–23. 10.1111/mec.13339

Hartl, D. L., Moriyama, E. N., & Sawyer, S. A. (1994). Selection intensity for codon bias. Genetics, 138(1), 227–234. 10.1093/genetics/138.1.227

Hellsten, U., Wright, K. M., Jenkins, J., Shu, S., Yuan, Y., Wessler, S. R., Schmutz, J., Willis, J. H., & Rokhsar, D. S. (2013). Fine-scale variation in meiotic recombination in *Mimulus* inferred from population shotgun sequencing. Proceedings of the National Academy of Sciences, 110(48), 19478–19482. 10.1073/pnas.1319032110

Hershberg, R., & Petrov, D. A. (2008). Selection on Codon Bias. Annual Review of Genetics, 42(1), 287–299. 10.1146/annurev.genet.42.110807.091442

Hill, W. G., & Robertson, A. (1966). The effect of linkage on limits to artificial selection. Genetical Research, 8(3), 269–294.

Hollister, J. D., Ross-Ibarra, J., & Gaut, B. S. (2010). Indel-Associated Mutation Rate Varies with Mating System in Flowering Plants. Molecular Biology and Evolution, 27(2), 409–416. 10.1093/molbev/msp249

Hsu, Y.-M., Falque, M., & Martin, O. C. (2022). Quantitative modelling of fine-scale variations in the Arabidopsis thaliana crossover landscape. Quantitative Plant Biology, 3, e3. 10.1017/qpb.2021.17

Ikemura, T. (1981). Correlation between the abundance of Escherichia coli transfer RNAs and the occurrence of the respective codons in its protein genes: A proposal for a synonymous codon choice that is optimal for the E. coli translational system. Journal of Molecular Biology, 151(3), 389–409. 10.1016/0022-2836(81)90003-6

Khandia, R., Saeed, Mohd., Alharbi, A. M., Ashraf, G. Md., Greig, N. H., & Kamal, M. A. (2022). Codon Usage Bias Correlates With Gene Length in Neurodegeneration Associated Genes. Frontiers in Neuroscience, 16, 895607. 10.3389/fnins.2022.895607

Koonin, E. V., & Novozhilov, A. S. (2009). Origin and evolution of the genetic code: The universal enigma. IUBMB Life, 61(2), 99–111. 10.1002/iub.146

Landerer, C., Cope, A., Zaretzki, R., & Gilchrist, M. A. (2018). AnaCoDa: Analyzing codon data with Bayesian mixture models. Bioinformatics, 34(14), 2496–2498. 10.1093/bioinformatics/bty138

Li, H., & Durbin, R. (2009). Fast and accurate short read alignment with Burrows–Wheeler transform. Bioinformatics, 25(14), 1754–1760. 10.1093/bioinformatics/btp324

Lovell, J. T., Walstead, R., Lawrence, A., Stark‐Dykema, E., Farnitano, M. C., Harder, A., Brůna, T., Barry, K., Goodstein, D., Jenkins, J., Lipzen, A., Boston, L., Webber, J., Chovatia, M., Eichenberger, J., Talag, J., Grimwood, J., Schmutz, J., Kelly, J. K., … Willis, J. H. (2025). Comparative Analyses of Four Reference Genomes Reveal Exceptional Diversity and Weak Linked Selection in the Yellow Monkeyflower (*Mimulus guttatus*) Complex. Molecular Ecology Resources, 25(8), e70012. 10.1111/1755-0998.70012

Lynch, M. (2010). Evolution of the mutation rate. Trends in Genetics, 26(8), 345–352. 10.1016/j.tig.2010.05.003

Lynch, M., Ackerman, M. S., Gout, J.-F., Long, H., Sung, W., Thomas, W. K., & Foster, P. L. (2016). Genetic drift, selection and the evolution of the mutation rate. Nature Reviews Genetics, 17(11), 704–714.

Machado, H. E., Lawrie, D. S., & Petrov, D. A. (2020). Pervasive Strong Selection at the Level of Codon Usage Bias in Drosophila melanogaster. Genetics, 214(2), 511–528. 10.1534/genetics.119.302542

Madrigal-Roca, L. J., Veltsos, P., & Kelly, J. K. (2025). Salmon Streamer: A Comprehensive Pipeline for RNA-seq to QTL Analysis (Version 1.0.0) [Computer software]. Zenodo. 10.5281/zenodo.XXXXXXX

McCandlish, D. M., & Plotkin, J. B. (2016). Transcriptional errors and the drift barrier. Proceedings of the National Academy of Sciences, 113(12), 3136–3138. 10.1073/pnas.1601785113

McDonald, J. H., & Kreitman, M. (1991). Adaptive protein evolution at the Adh locus in Drosophila. Nature, 351(6328), 652–654. 10.1038/351652a0

McVEAN, G. A. T., & Charlesworth, B. (1999). A population genetic model for the evolution of synonymous codon usage: Patterns and predictions. Genetical Research, 74(2), 145–158. 10.1017/S0016672399003912

Murphy, F. V., & Ramakrishnan, V. (2004). Structure of a purine-purine wobble base pair in the decoding center of the ribosome. Nature Structural & Molecular Biology, 11(12), 1251–1252. 10.1038/nsmb866

Niehrs, C., & Luke, B. (2020). Regulatory R-loops as facilitators of gene expression and genome stability. Nature Reviews Molecular Cell Biology, 21(3), 167–178. 10.1038/s41580-019-0206-3

Nielsen, R. (2001). Statistical tests of selective neutrality in the age of genomics. Heredity, 86(6), 641–647. 10.1046/j.1365-2540.2001.00895.x

Okagaki, R. J., Dukowic-Schulze, S., Eggleston, W. B., & Muehlbauer, G. J. (2018). A Critical Assessment of 60 Years of Maize Intragenic Recombination. Frontiers in Plant Science, 9, 1560. 10.3389/fpls.2018.01560

Pál, C., Papp, B., & Hurst, L. D. (2001). Highly Expressed Genes in Yeast Evolve Slowly. Genetics, 158(2), 927–931. 10.1093/genetics/158.2.927

Parvathy, S. T., Udayasuriyan, V., & Bhadana, V. (2022). Codon usage bias. Molecular Biology Reports, 49(1), 539–565. 10.1007/s11033-021-06749-4

Patro, R., Duggal, G., Love, M. I., Irizarry, R. A., & Kingsford, C. (2017). Salmon provides fast and bias-aware quantification of transcript expression. Nature Methods, 14(4), 417–419. 10.1038/nmeth.4197

Perriere, G. (2002). Use and misuse of correspondence analysis in codon usage studies. Nucleic Acids Research, 30(20), 4548–4555. 10.1093/nar/gkf565

Picard toolkit. (2019). In Broad Institute, GitHub repository. Broad Institute. https://broadinstitute.github.io/picard/

Polak, P., & Arndt, P. F. (2008). Transcription induces strand-specific mutations at the 5′ end of human genes. Genome Research, 18(8), 1216–1223. 10.1101/gr.076570.108

R Core Team. (2024). R: A Language and Environment for Statistical Computing. R Foundation for Statistical Computing. https://www.R-project.org/

Sharp, P. M., & Li, W.-H. (1987). The codon adaptation index-a measure of directional synonymous codon usage bias, and its potential applications. Nucleic Acids Research, 15(3), 1281–1295. 10.1093/nar/15.3.1281

Sharp, P. M., Tuohy, T. M. F., & Mosurski, K. R. (1986). Codon usage in yeast: Cluster analysis clearly differentiates highly and lowly expressed genes. Nucleic Acids Research, 14(13), 5125–5143. 10.1093/nar/14.13.5125

Su, X. A., & Freudenreich, C. H. (2017). Cytosine deamination and base excision repair cause R-loop–induced CAG repeat fragility and instability in *Saccharomyces cerevisiae*. Proceedings of the National Academy of Sciences, 114(40). 10.1073/pnas.1711283114

Sun, M., & Zhang, J. (2022). Preferred synonymous codons are translated more accurately: Proteomic evidence, among-species variation, and mechanistic basis. Science Advances, 8(27), eabl9812. 10.1126/sciadv.abl9812

Sun, X., Yang, Q., & Xia, X. (2013). An Improved Implementation of Effective Number of Codons (Nc). Molecular Biology and Evolution, 30(1), 191–196. 10.1093/molbev/mss201

Troth, A., Puzey, J. R., Kim, R. S., Willis, J. H., & Kelly, J. K. (2018). Selective trade-offs maintain alleles underpinning complex trait variation in plants. Science, 361(6401), 475–478. 10.1126/science.aat5760

Trucchi, E., Massa, P., Giannelli, F., Latrille, T., Fernandes, F. A. N., Ancona, L., Stenseth, N. C., Ferrer Obiol, J., Paris, J., Bertorelle, G., & Le Bohec, C. (2023). Gene expression is the main driver of purifying selection in large penguin populations. Evolutionary Biology. 10.1101/2023.08.08.552445

Vogl, C. (2014). Estimating the scaled mutation rate and mutation bias with site frequency data. Theoretical Population Biology, 98, 19–27. 10.1016/j.tpb.2014.10.002

Wang, H.-C., & Hickey, D. A. (2007). Rapid divergence of codon usage patterns within the rice genome. BMC Evolutionary Biology, 7(Suppl 1), S6. 10.1186/1471-2148-7-S1-S6

Wijnker, E., Velikkakam James, G., Ding, J., Becker, F., Klasen, J. R., Rawat, V., Rowan, B. A., De Jong, D. F., De Snoo, C. B., Zapata, L., Huettel, B., De Jong, H., Ossowski, S., Weigel, D., Koornneef, M., Keurentjes, J. J., & Schneeberger, K. (2013). The genomic landscape of meiotic crossovers and gene conversions in Arabidopsis thaliana. eLife, 2, e01426. 10.7554/eLife.01426

Wood, S. N. (2004). Stable and efficient multiple smoothing parameter estimation for generalized additive models. Journal of the American Statistical Association, 99(467), 673–686.

Wood, S. N. (2011). Fast stable restricted maximum likelihood and marginal likelihood estimation of semiparametric generalized linear models. Journal of the Royal Statistical Society (B), 73(1), 3–36.

Wood, S. N., N., Pya, & S”afken, B. (2016). Smoothing parameter and model selection for general smooth models (with discussion). Journal of the American Statistical Association, 111, 1548–1575.

Wright, F. (1990). The ‘effective number of codons’ used in a gene. Gene, 87(1), 23–29. 10.1016/0378-1119(90)90491-9

Wright, S. (1931). EVOLUTION IN MENDELIAN POPULATIONS. Genetics, 16(2), 97–159. 10.1093/genetics/16.2.97

Zhang, J., & Yang, J.-R. (2015). Determinants of the rate of protein sequence evolution. Nature Reviews Genetics, 16(7), 409–420. 10.1038/nrg3950

Zhang, Z., Li, J., Cui, P., Ding, F., Li, A., Townsend, J. P., & Yu, J. (2012). Codon Deviation Coefficient: A novel measure for estimating codon usage bias and its statistical significance. BMC Bioinformatics, 13(1), 43. 10.1186/1471-2105-13-43

