## Supplemental materials for "The dynamics of “silent” variation in *Mimulus guttatus*: Codon usage bias and linked selection"

### **Supplementary results**

ROC_SEMPPR fitting

Independent chains fit under the naive model failed to converge to a unique solution, yielding Gelman-Rubin convergence diagnostics ($\hat{R}$) consistently greater than 1.1. This stems from a lack of identifiability: without informative priors, the model cannot distinguish between a regime where mutation favors AT (counteracted by selection for GC) and the inverse scenario. Of the models where the mutational bias ($\Delta M$) was fixed to intronic or intergenic estimates without expression information, only the one based on introns converged successfully. Nevertheless, the estimated $\Phi$ values were negatively correlated with the empirical expression values, suggesting that the simultaneous estimation of selection and the mean protein production rate lead to misleading results ($\rho_{\Delta M_{introns}} = -0.16, df = 22{,}568, p < 2.2 \cdot 10^{-16}$; $\rho_{\Delta M_{intergenic}} = -0.13, df = 22{,}568, p < 2.2 \cdot 10^{-16}$).

Finally, from the models where empirical expression data was used, given that both achieved convergence, we selected the one where mutational bias was set based on intronic sequences. As shown in Table 1, introns exhibit a stronger, strand-specific T-bias compared to intergenic regions; because introns are transcribed and reside in open chromatin, their mutational profile more accurately reflects the load experienced by CDS blocks. That is why we based our downstream inferences based on the dM-fixed-with_phi model, which estimates $\Delta\eta$ conditional in the observed values of gene expression and the intronic mutational bias. With exception of the codon CTT for Lysine, the differences in cost estimated from our best ROC-SEMPPR model are statistically significant, with no CI at 95% including 0 (Fig. S4).

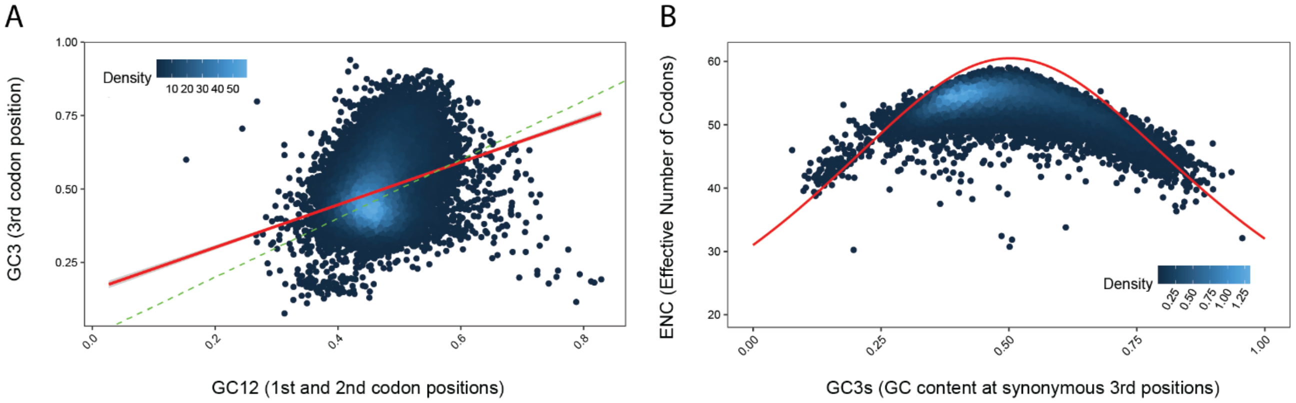


Supplementary Figure 1: General patterns of Codon Usage Bias in *Mimulus guttatus* from Iron Mountain, Oregon, USA. **A)** Neutrality plot of GC12 and GC3, where the green dashed line represent the expectation under mutation only. A deviation towards slope 0 is a signature of selection acting to shape the codon landscape. The red line is the empirical regression model based on CUB in the yellow monkey flower. **B)** ENC plot showing the ENC expectation as a function of GC3s content as a red line. Notice that GC3s considers only 18 of the 20 amino acid families, the ones that are coded at least for two codons. Each dot represents a gene, and those under the curve are pushed presumably by selective forces.

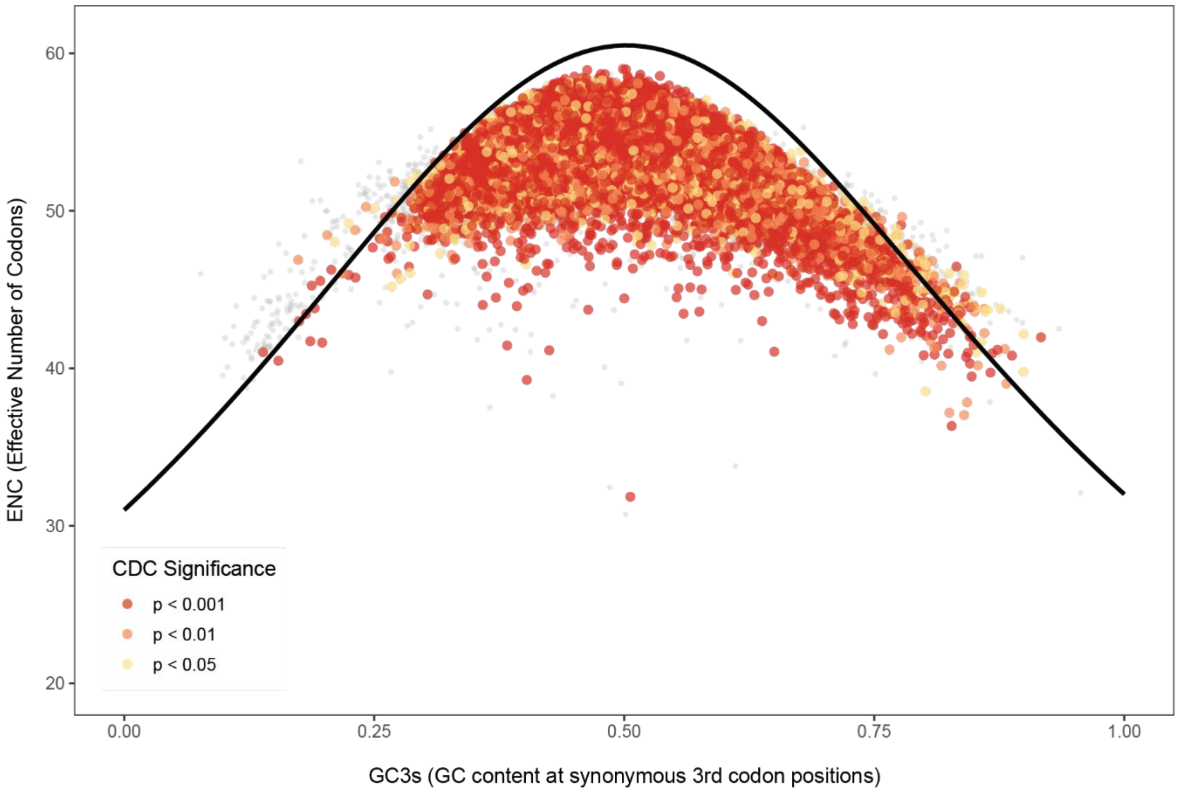


Supplementary Figure 2: ENC plot showing the ENC expectation as a function of GC3s content as a black line. Notice that GC3s considers only 18 of the 20 amino acid families, the ones that are coded at least for two codons. Each dot represent a gene, which have been colored as a function of significance levels derived from the CDC calculations.

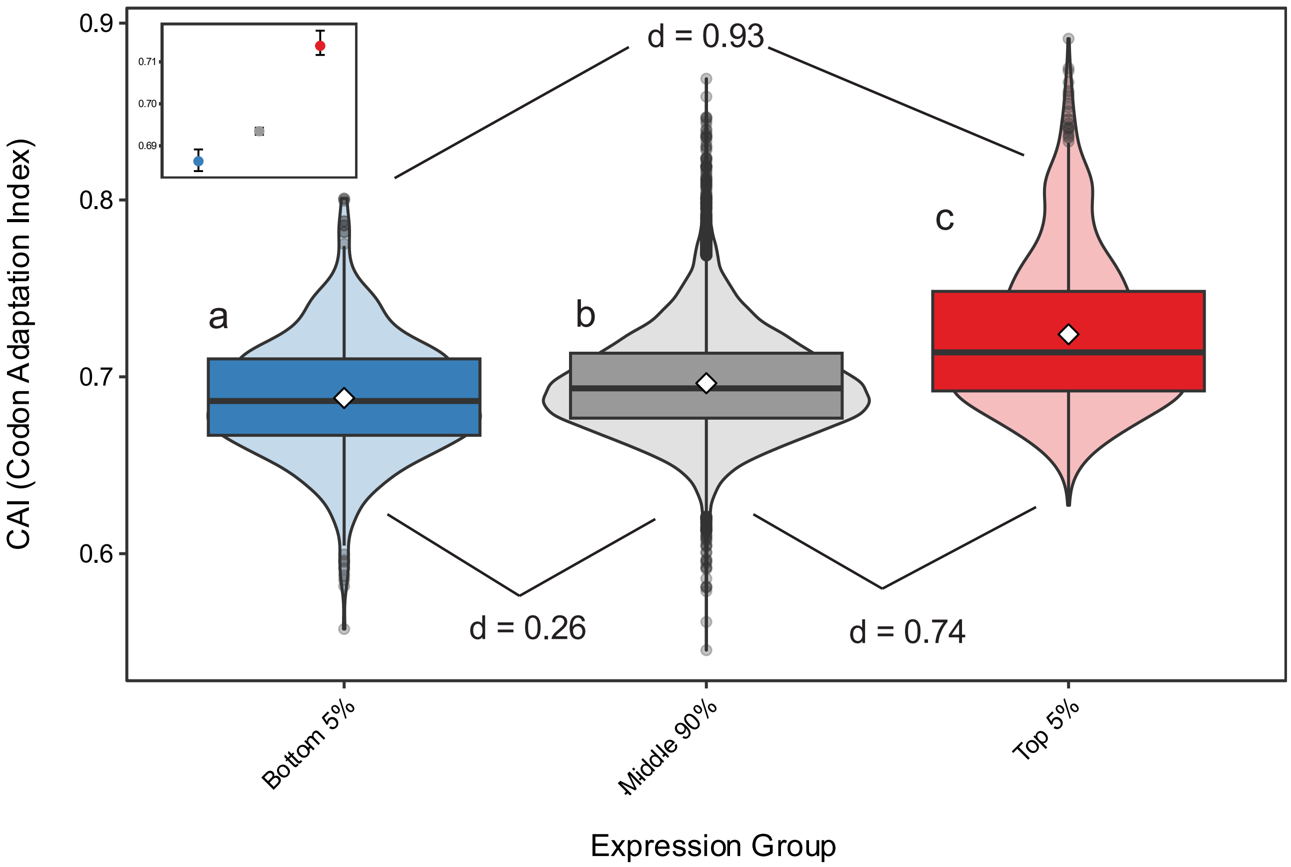


Supplementary Figure 3: Codon Adaptation Index (CAI) and its relationship with expression in the yellow monkey flower, *Mimulus guttatus*, from the natural population Iron Mountain, Oregon, USA. For each expression group, the violin represents the empirical CAI residual distribution, the box represents the inter quartile range, diamonds represent the mean and horizontal line the median. Lowercase letter next to each box denominates the results from the Dunn test (FDR < 0.05). d values represent the Cohen's d estimate of effect size, where |d| < 0.2 = negligible, 0.2-0.5 = small, 0.5-0.8 = medium, > 0.8 = large. Inset represents the median and its 95% confidence interval.

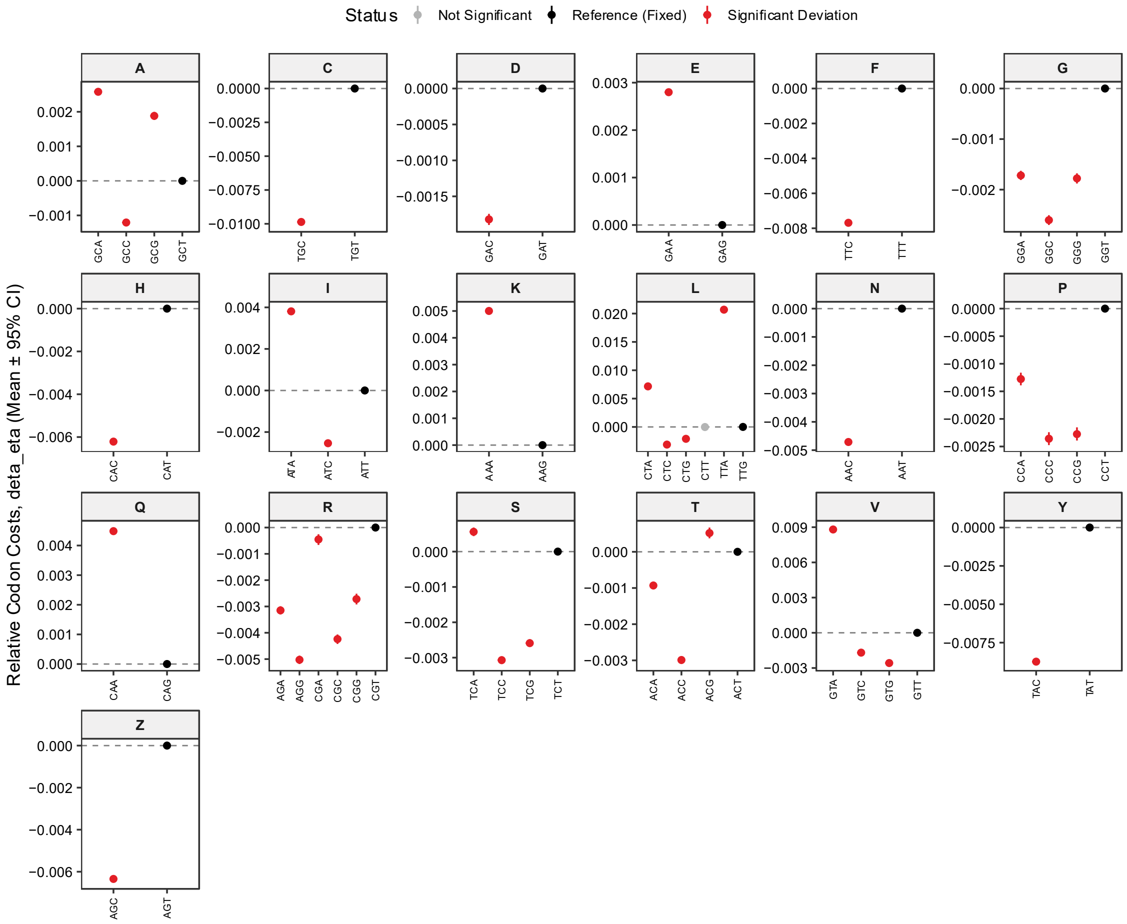


Supplementary Figure 4: Codon inefficiencies relative to a reference (black dots) estimated from AnaCoDa models for *Mimulus guttatus* from Iron Mountain, Oregon, USA. Generally, GC-ending codons seem to be less costly from the translational perspective, with exception of the ACG codon in Tyrosine.

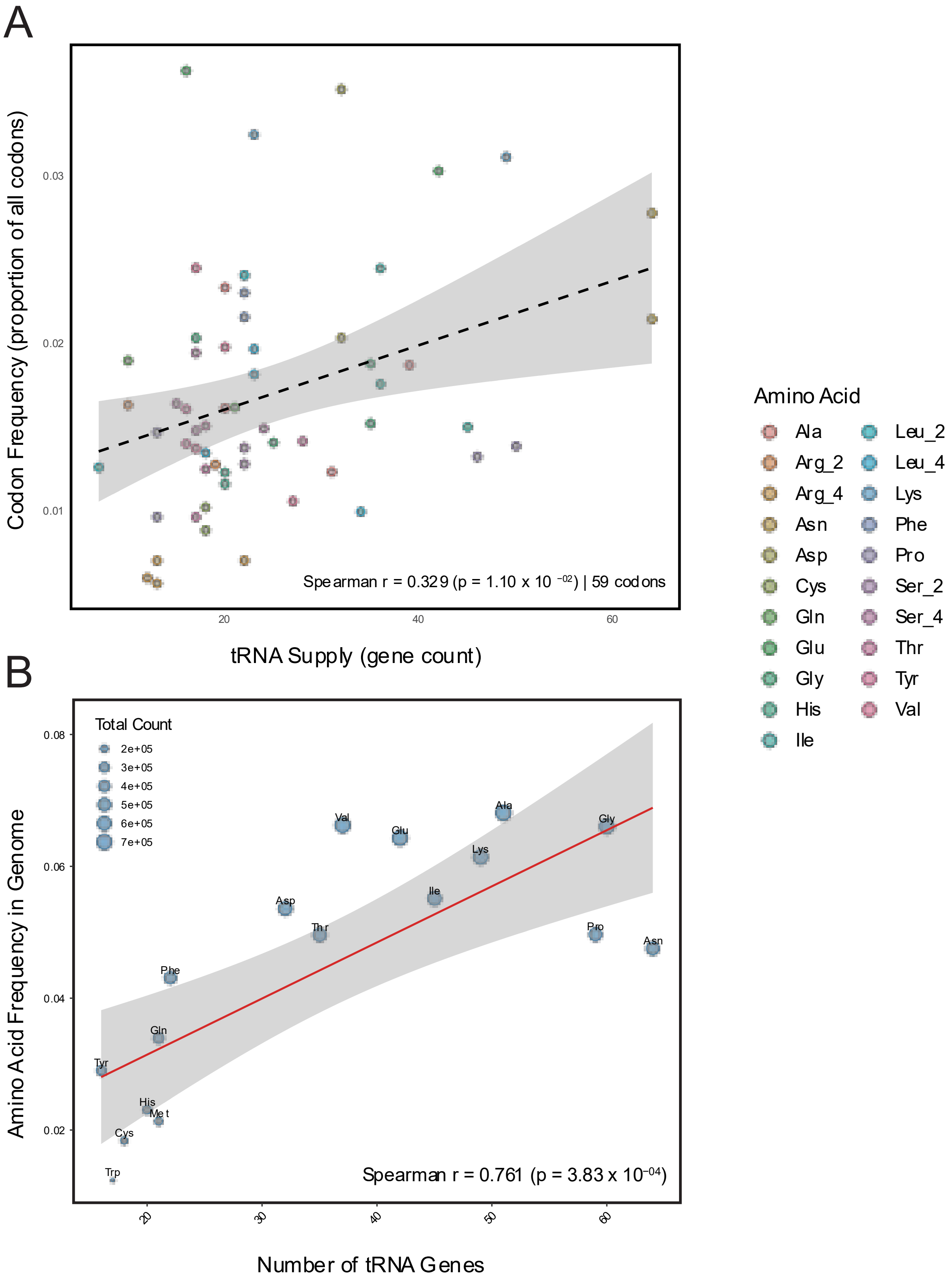


Supplementary Figure 5: Co-evolution of tRNA gene supply and genomic composition in *Mimulus guttatus*. **(A)** Relationship between synonymous codon frequency (expressed as a proportion of all codons) and tRNA gene copy number. A positive correlation $(Spearman\rho = 0.329, p = 1.10 \times 10^{-2})$ suggests that genomic codon usage has evolved to match the available tRNA pool to optimize translational efficiency. **(B)** Correlation between total amino acid frequency in the *M.*

*guttatus* genome and the number of corresponding tRNA genes. The high correlation coefficient ($Spearman\rho = 0.761, p = 3.83 \times 10^{-4}$) demonstrates that the overall amino acid composition of the proteome is strongly coupled with the translational capacity of the cell. Points are color-coded by amino acid; the dashed line represents the linear regression fit with the 95% confidence interval shaded in gray.

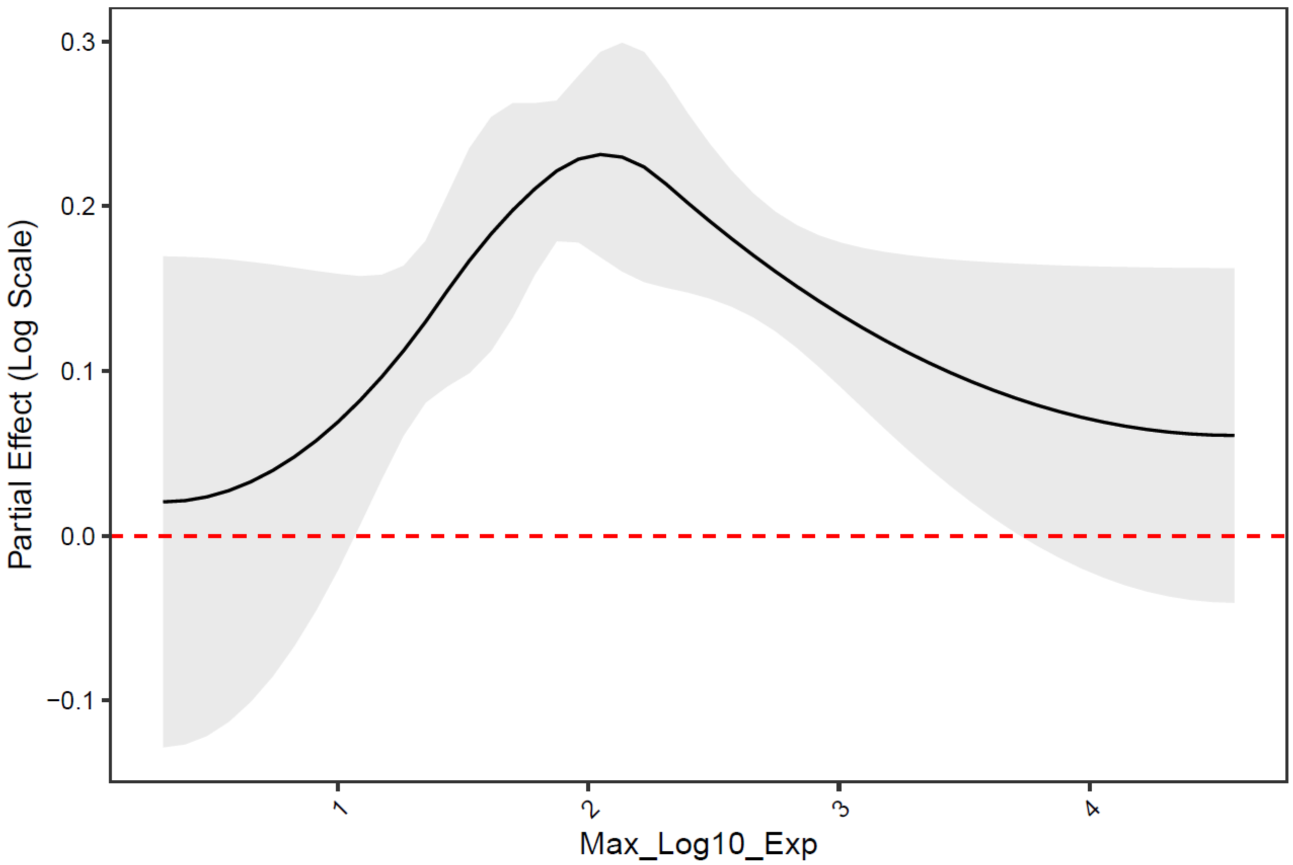


Supplementary Figure 6: Partial Effect of Expression Intensity on Preferred Codon Frequency. This plot illustrates the non-linear relationship between maximum expression intensity ($log_{10}$ scale) and the mean frequency of preferred codons, while controlling for other model predictors such as expression breadth and gene length. The solid black line represents the estimated partial effect, showing a predominantly positive influence that peaks at intermediate expression levels ($\approx 10^2$) before exhibiting a gradual decline at the highest intensities. The grey shaded region denotes the 95% confidence interval; where this region remains entirely above the red dashed null line (0.0), the effect of expression on preferred codon frequency is statistically significant. This trend suggests that while increased expression intensity generally drives higher usage of preferred codons to optimize translational efficiency, the strength of this selective pressure may reach a plateau or vary in extremely highly expressed genes.

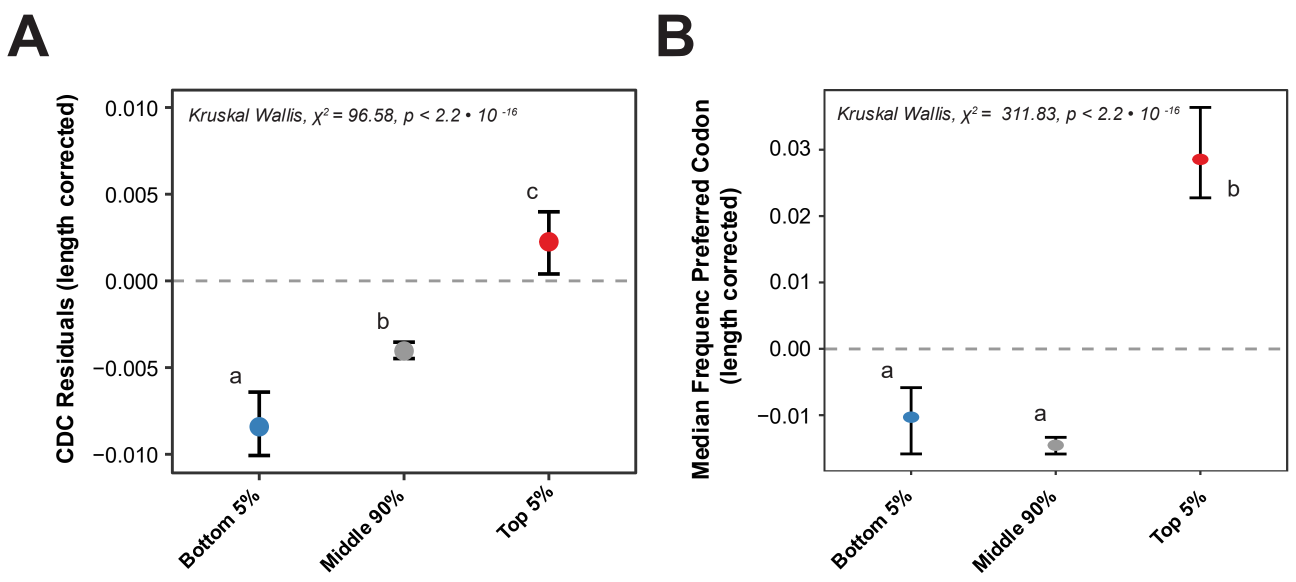


Supplementary Figure 7: Codon Deviation Coefficient Residuals (CDC Residuals) and its relationship with expression in the yellow monkey flower, *Mimulus guttatus*, from the natural population Iron Mountain, Oregon, USA. **A)** De-trended CDC metric after controlling for gene length using a Generalized Additive Model (beta family, logit link function). For each expression group, dot represent the median and the whiskers the 95% CI. Lowercase letter next to each box denominates the results from the Dunn test (FDR < 0.05). **B)** De-trended mean frequency of the preferred allele (*p*) after controlling for gene length using a Generalized Additive Model (beta family, logit link function). For each expression group, the dot represents the median and the whiskers the 95% CI. Lowercase letter next to each box denominates the results from the Dunn test (FDR < 0.05). Highly and lowly expressed genes in a bi-dimensional space defined by dimensions 1 and 3 of a correspondence analysis.

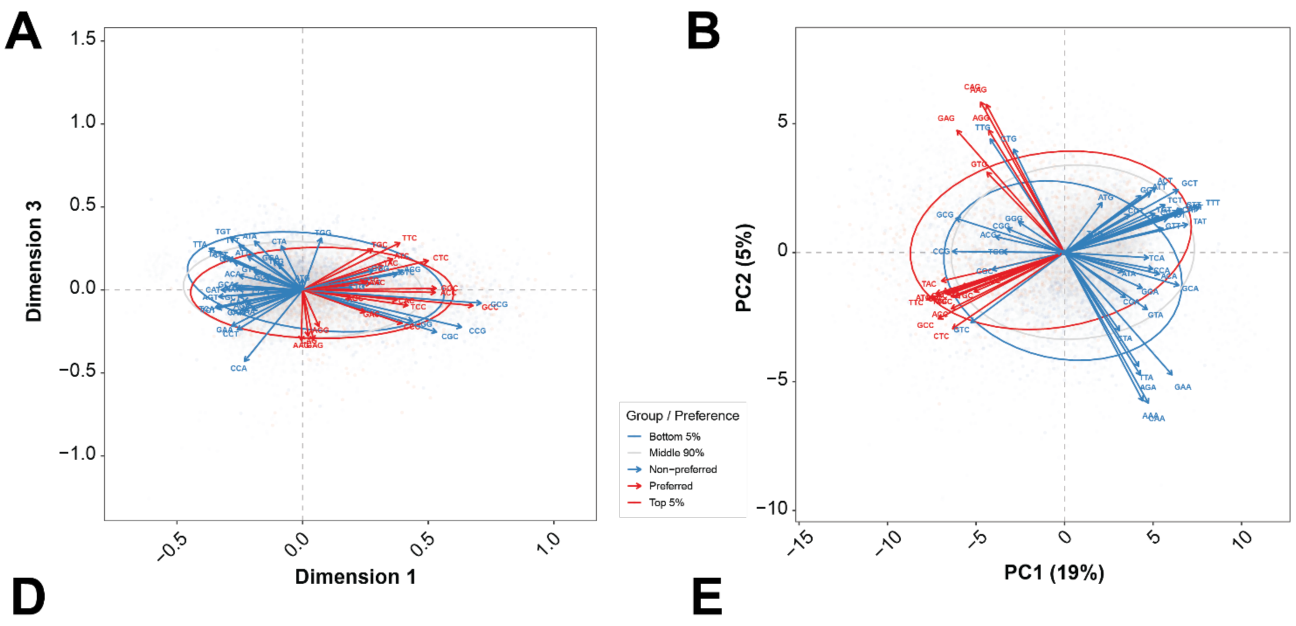


Supplementary Figure 8: Highly and lowly expressed genes in a bi-dimensional space defined by dimensions 1 and 3 of a correspondence analysis (A) based on codon counts and the two first principal components from a PCA based on the RSCU scores (B) for *Mimulus guttatus* from iron Mountain, Oregon.

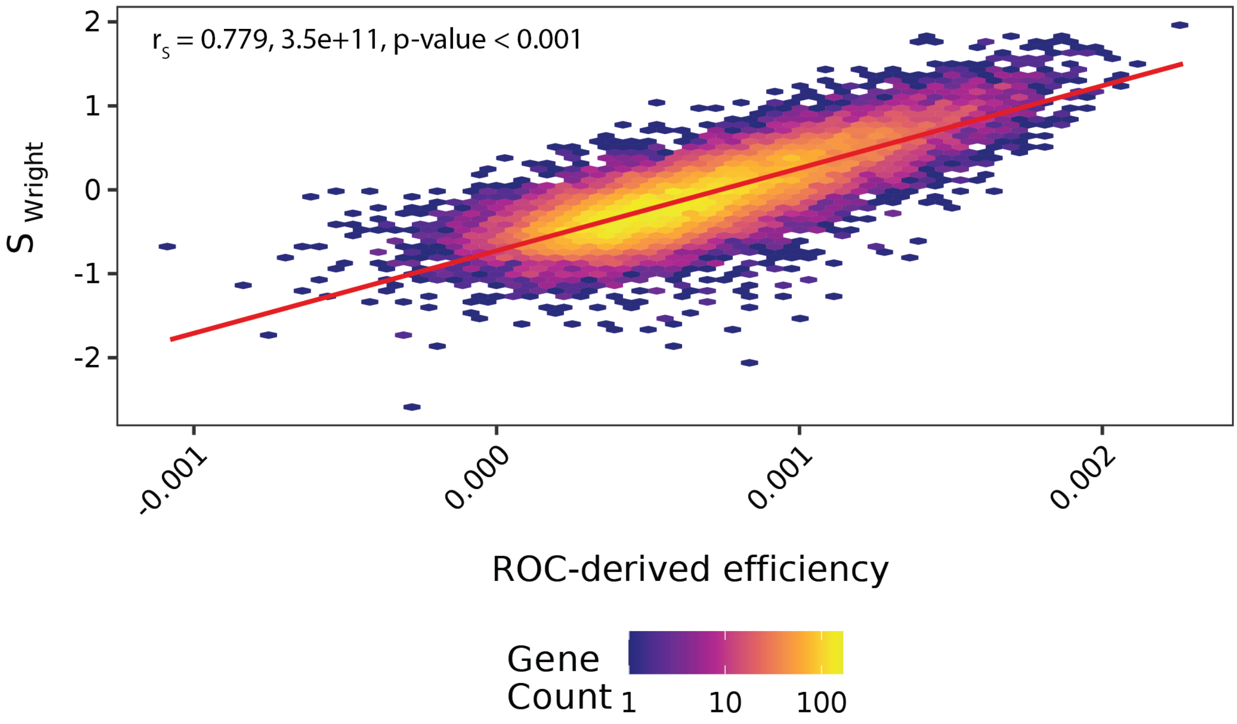


Supplementary Figure 9: Relationship between ROC-derived efficiencies and the predictions of Wright equation after transforming the nucleotide composition at 4-fold sites into a two-allele system (preferred/unpreferred bases).

Table S1: G-based heterogeneity test for CUB across amino acid families.

| Amino acid | $G_{het}$ | *df* | $p_{value}(adj)$ |
|---|---|---|---|
| Ala | 184508.5 | 75462 | $< 2.2 \cdot 10^{-16}$ |
| Arg_2 | 36294.9 | 24694 | $< 2.2 \cdot 10^{-16}$ |
| Arg_4 | 114949.0 | 73755 | $< 2.2 \cdot 10^{-16}$ |
| Asn | 52707.5 | 25076 | $< 2.2 \cdot 10^{-16}$ |
| Asp | 48659.8 | 25042 | $< 2.2 \cdot 10^{-16}$ |
| Cys | 39132.7 | 23989 | $< 2.2 \cdot 10^{-16}$ |
| Gln | 44747.2 | 24941 | $< 2.2 \cdot 10^{-16}$ |
| Glu | 47117.1 | 25072 | $< 2.2 \cdot 10^{-16}$ |
| Gly | 132528.6 | 75435 | $< 2.2 \cdot 10^{-16}$ |
| His | 47086.2 | 24482 | $< 2.2 \cdot 10^{-16}$ |
| Ile | 86330.2 | 50274 | $< 2.2 \cdot 10^{-16}$ |
| Leu_2 | 39316.3 | 24827 | $< 2.2 \cdot 10^{-16}$ |
| Leu_4 | 138313.0 | 75372 | $< 2.2 \cdot 10^{-16}$ |
| Lys | 42769.5 | 25118 | $< 2.2 \cdot 10^{-16}$ |
| Phe | 56752.8 | 25068 | $< 2.2 \cdot 10^{-16}$ |
| Pro | 148909.2 | 75225 | $< 2.2 \cdot 10^{-16}$ |
| Ser_2 | 42262.9 | 24717 | $< 2.2 \cdot 10^{-16}$ |
| Ser_4 | 160356.7 | 75372 | $< 2.2 \cdot 10^{-16}$ |
| Thr | 139580.4 | 75384 | $< 2.2 \cdot 10^{-16}$ |
| Tyr | 49820.7 | 24764 | $< 2.2 \cdot 10^{-16}$ |
| Val | 123393.1 | 75483 | $< 2.2 \cdot 10^{-16}$ |

Table S2: GO enrichment results from genes where $S_{Wright} > 1$ in *Mimulus guttatus* from Iron Mountain, Oregon.

| Term ID | Term name | p value |
|---|---|---|
| GO:0140678 | molecular function inhibitor activity | 3.37E-17 |
| GO:0004857 | enzyme inhibitor activity | 6.09E-17 |
| GO:0030234 | enzyme regulator activity | 2.57E-15 |
| GO:0003735 | structural constituent of ribosome | 3.22E-13 |
| GO:0005198 | structural molecule activity | 4.42E-11 |
| GO:0016747 | acyltransferase activity, transferring groups other than amino-acyl groups | 1.23E-08 |
| GO:0016757 | glycosyltransferase activity | 1.51E-08 |
| GO:0016758 | hexosyltransferase activity | 1.10E-07 |
| GO:0016746 | acyltransferase activity | 1.10E-07 |
| GO:0071555 | cell wall organization | 1.33E-07 |
| GO:0030599 | pectinesterase activity | 6.22E-07 |
| GO:0042545 | cell wall modification | 6.22E-07 |
| GO:0006412 | translation | 6.22E-07 |
| GO:0004097 | catechol oxidase activity | 6.22E-07 |
| GO:0045229 | external encapsulating structure organization | 2.90E-06 |
| GO:0004497 | monooxygenase activity | 2.00E-05 |
| GO:0045735 | nutrient reservoir activity | 2.19E-05 |
| GO:0052689 | carboxylic ester hydrolase activity | 5.35E-05 |
| GO:0009733 | response to auxin | 5.80E-05 |
| GO:0006817 | phosphate ion transport | 7.17E-05 |
| GO:0005315 | phosphate transmembrane transporter activity | 7.17E-05 |
| GO:0046148 | pigment biosynthetic process | 0.000246 |
| GO:0015035 | protein-disulfide reductase activity | 0.000283 |
| GO:0071554 | cell wall organization or biogenesis | 0.000307 |
| GO:0008519 | ammonium channel activity | 0.000438 |
| GO:0072488 | ammonium transmembrane transport | 0.000438 |
| GO:0009055 | electron transfer activity | 0.000492 |
| GO:0046982 | protein heterodimerization activity | 0.000647 |
| GO:0042440 | pigment metabolic process | 0.001064 |
| GO:0042221 | response to chemical | 0.001214 |
| GO:0016667 | oxidoreductase activity, acting on a sulfur group of donors | 0.001342 |
| GO:0000786 | nucleosome | 0.003274 |
| GO:0032993 | protein-DNA complex | 0.003274 |
| GO:0009579 | thylakoid | 0.005416 |
| GO:0005506 | iron ion binding | 0.005964 |
| GO:0006814 | sodium ion transport | 0.006095 |
| GO:0015036 | disulfide oxidoreductase activity | 0.006095 |
| GO:0016705 | oxidoreductase activity, acting on paired donors, with incorporation or reduction of molecular oxygen | 0.006194 |
| GO:0015698 | inorganic anion transport | 0.007123 |
| GO:0008610 | lipid biosynthetic process | 0.008071 |
| GO:1902680 | positive regulation of RNA biosynthetic process | 0.008878 |
| GO:0045893 | positive regulation of DNA-templated transcription | 0.008878 |
| GO:0000785 | chromatin | 0.009577 |
| GO:0009521 | photosystem | 0.009824 |

| | | |
|---|---|---|
| GO:0034357 | photosynthetic membrane | 0.009824 |
| GO:0046906 | tetrapyrrole binding | 0.012186 |
| GO:0031122 | cytoplasmic microtubule organization | 0.012314 |
| GO:0009538 | photosystem I reaction center | 0.012314 |
| GO:0000930 | gamma-tubulin complex | 0.012314 |
| GO:0015979 | photosynthesis | 0.012808 |
| GO:0020037 | heme binding | 0.013045 |
| GO:0006633 | fatty acid biosynthetic process | 0.013614 |
| GO:0009765 | photosynthesis, light harvesting | 0.0154 |
| GO:0009725 | response to hormone | 0.015686 |
| GO:0009719 | response to endogenous stimulus | 0.015686 |
| GO:0072330 | monocarboxylic acid biosynthetic process | 0.015939 |
| GO:0005200 | structural constituent of cytoskeleton | 0.015939 |
| GO:0008150 | biological_process | 0.017229 |
| GO:0046785 | microtubule polymerization | 0.022964 |
| GO:0005815 | microtubule organizing center | 0.022964 |
| GO:0007020 | microtubule nucleation | 0.022964 |
| GO:0008152 | metabolic process | 0.025815 |
| GO:0009522 | photosystem I | 0.025815 |
| GO:0009058 | biosynthetic process | 0.029894 |
| GO:0016740 | transferase activity | 0.038069 |
| GO:0098660 | inorganic ion transmembrane transport | 0.039573 |
| GO:0140352 | export from cell | 0.039573 |
| GO:0046903 | secretion | 0.039573 |
| GO:0032940 | secretion by cell | 0.039573 |
| GO:0042592 | homeostatic process | 0.047933 |
| GO:0016491 | oxidoreductase activity | 0.048577 |

Table S3: Top genes from *Mimulus guttatus* as a function of the translational load (coefficient, $L_{ROC}$) and their best hit in *Arabidopsis thaliana.*

| Gene_name | L_ROC | Best hit *Arabidopsis* name | Best hit *Arabidopsis* definition |
|---|---|---|---|
| MgIM767.12G162800 | 2.692620053 | AT5G17920 | Cobalamin-independent synthase family protein |
| MgIM767.08G165400 | 2.422288833 | AT5G56010 | heat shock protein 81-3 |
| MgIM767.10G141000 | 2.346082934 | AT2G18960 | H(+)-ATPase 1 |
| MgIM767.13G003200 | 2.186252096 | AT2G05070 | photosystem II light harvesting complex gene 2.2 |
| MgIM767.12G169900 | 2.142013747 | AT5G02500 | heat shock cognate protein 70-1 |
| MgIM767.04G191000 | 2.103040714 | AT2G27800 | Tetratricopeptide repeat (TPR)-like superfamily protein |
| MgIM767.08G160300 | 2.06435118 | NA | NA |
| MgIM767.08G042100 | 2.057106599 | AT3G50820 | photosystem II subunit O-2 |
| MgIM767.06G077600 | 2.047007607 | AT3G12580 | heat shock protein 70 |
| MgIM767.14G110400 | 2.028199024 | NA | NA |
| MgIM767.04G117800 | 1.957301639 | AT5G50920 | CLPC homologue 1 |
| MgIM767.12G087300 | 1.818571388 | AT3G25570 | Adenosylmethionine decarboxylase family protein |
| MgIM767.08G117700 | 1.638631035 | AT1G06680 | photosystem II subunit P-1 |

| | | | |
|---|---|---|---|
| MgIM767.06G100000 | 1.629526889 | AT5G42020 | Heat shock protein 70 (Hsp 70) family protein |
| MgIM767.02G150900 | 1.581210633 | AT3G60750 | Transketolase |
| MgIM767.03G128800 | 1.528469113 | AT4G10340 | light harvesting complex of photosystem II 5 |
| MgIM767.06G075100 | 1.488247085 | AT2G39730 | rubisco activase |
| MgIM767.04G190700 | 1.423887742 | AT1G07940 | GTP binding Elongation factor Tu family protein |
| MgIM767.05G133800 | 1.413264279 | AT2G26080 | glycine decarboxylase P-protein 2 |
| MgIM767.04G058200 | 1.385714845 | AT3G08580 | ADP/ATP carrier 1 |
| MgIM767.09G049300 | 1.38482749 | AT5G38430 | Ribulose bisphosphate carboxylase (small chain) family protein |
| MgIM767.09G060900 | 1.374255249 | NA | NA |
| MgIM767.02G019400 | 1.361001962 | AT4G39080 | vacuolar proton ATPase A3 |
| MgIM767.14G051000 | 1.353001066 | AT1G55490 | chaperonin 60 beta |
| MgIM767.08G008200 | 1.349967135 | AT5G40450 | Regulator of bulb biogenesis1 |
| MgIM767.11G119700 | 1.318952854 | NA | NA |
| MgIM767.05G037900 | 1.308617498 | AT5G08690 | ATP synthase alpha/beta family protein |
| MgIM767.08G195100 | 1.297660294 | AT1G61520 | photosystem I light harvesting complex gene 3 |
| MgIM767.02G085500 | 1.267936026 | NA | NA |
| MgIM767.08G021800 | 1.267281404 | NA | NA |
| MgIM767.02G165700 | 1.258184481 | AT3G61430 | plasma membrane intrinsic protein 1A |
| MgIM767.12G121000 | 1.254460359 | AT1G60470 | galactinol synthase 4 |
| MgIM767.13G039000 | 1.24055061 | AT1G06430 | FTSH protease 8 |
| MgIM767.10G069000 | 1.230690552 | AT5G17920 | Cobalamin-independent synthase family protein |
| MgIM767.06G066100 | 1.200323554 | AT3G11130 | Clathrin, heavy chain |
| MgIM767.14G261200 | 1.199345088 | AT1G49760 | poly(A) binding protein 8 |
| MgIM767.11G130500 | 1.19683575 | NA | NA |
| MgIM767.03G066000 | 1.194437607 | NA | NA |
| MgIM767.02G017600 | 1.132281048 | AT4G38970 | fructose-bisphosphate aldolase 2 |
| MgIM767.04G139300 | 1.101842497 | AT5G38430 | Ribulose bisphosphate carboxylase (small chain) family protein |
| MgIM767.09G005200 | 1.066799047 | AT1G48410 | Stabilizer of iron transporter SufD / Polynucleotidyl transferase |
| MgIM767.13G018400 | 1.059183559 | NA | NA |
| MgIM767.07G076900 | 1.055628451 | AT3G16640 | translationally controlled tumor protein |
| MgIM767.13G156500 | 1.047509129 | AT1G55860 | ubiquitin-protein ligase 1 |
| MgIM767.10G139300 | 1.026201073 | AT4G30210 | P450 reductase 2 |
| MgIM767.10G166100 | 1.01196427 | AT5G13630 | magnesium-chelatase subunit chlH, chloroplast, putative / Mg-protoporphyrin IX chelatase, putative (CHLH) |
| MgIM767.08G158400 | 1.004938995 | AT1G79550 | phosphoglycerate kinase |
| MgIM767.04G172600 | 0.999345765 | AT4G24690 | ubiquitin-associated (UBA)/TS-N domain-containing protein / octicosapeptide/Phox/Bemp1 (PB1) do-main-containing protein |
| MgIM767.08G158300 | 0.974446008 | AT3G12780 | phosphoglycerate kinase 1 |
| MgIM767.01G088400 | 0.957224219 | NA | NA |

Table S4: Summary of biological replicates sequenced per tissue type for lines IM62 and IM767 of *Mimulus guttatus* from Iron Mountain, Oregon.

| Tissue Category | Tissue Type | IM62 (n=15) | IM767 (n=14) | Total (n=29) |
|---|---|---|---|---|
| **Vegetative** | Leaf | 4 | 4 | 8 |
| | Root(s) | 1 | 1 | 2 |
| | Stem | 2 | 1 | 3 |
| | Seedling | 0 | 1 | 1 |
| **Reproductive** | Bud | 2 | 1 | 3 |
| | Flower | 2 | 1 | 3 |
| | Ovary | 2 | 1 | 3 |
| | Pollen | 0 | 1 | 1 |
| **Mixed** | Pool | 2 | 3 | 5 |
